# Stress-driven emergence of heritable non-genetic drug resistance

**DOI:** 10.1101/2024.12.20.629823

**Authors:** Jinglin L. Xie, Sifei Yin, Theodore S. Yang, Kiran Chandrasekher, Luke Hanson, Sang Hu Kim, Lucas Esqueda, Catherine A. Hogan, Niaz Banaei, June L. Round, Kyla S. Ost, Judith Berman, Daniel F. Jarosz

## Abstract

Drug resistance is the chief cause of treatment failure for therapies targeting chronic and infectious diseases. Whether the emergence of resistance is accelerated by environmental exposure to low levels of therapeutics remains controversial. Here, we report a non-genetic mechanism of stress adaptation that promotes heritable resistance to the widely used antifungal drug fluconazole. In the human fungal pathogen *Candida albicans,* transient exposure to subtherapeutic fluconazole doses induces a protective response that we term para-resistance. Like conventional resistance mechanisms, para-resistance is heritable. However, it does not arise from genetic mutations and can revert spontaneously. Systematic analyses of para-resistant isolates suggest that its key regulators include the stress-activated MAP kinase Hog1, the histone deacetylase subunit Snt1, the chromatin regulator Rap1, and the Sko1 transcriptional factor. Notably, molecules that disrupt biomolecular condensation and prion propagation – crucial for the inheritance of protein assemblies – block the induction of para-resistance, whereas inhibiting histone deacetylases facilitates its induction. We find that para-resistance is common in clinical isolates and, remarkably, passage through the mammalian gut triggers its acquisition, compromising fluconazole’s therapeutic efficacy. Our work defines a pervasive, prion-like epigenetic mechanism of stress adaptation and highlights potential strategies to mitigate the rapid emergence of drug resistance.

## Introduction

Drug resistance is a chief cause of treatment failure in oncology and infectious disease.^1,2^ Antibacterial resistance alone kills >1 million annually.^1^ In agriculture, the cost of crop loss due to pesticide resistance is estimated to be 1.5 billion USD per year.^3^ Beyond the risk of eroding some of the greatest advances in human health and agricultural productivity achieved over the past century, the prodigious emergence of drug resistance also poses a fascinating evolutionary puzzle. Forward and reverse genetics have greatly expanded our understanding of how mutations contribute to drug resistance.^4–6^ However, emerging evidence has sparked interest in whether non-genetic mechanisms may act as a hidden force, facilitating rapid adaptation to drug-induced stress.^7–9^ One line of evidence centers on the observation that many infections fail to respond to treatment despite exhibiting population-wide drug susceptibility.^10–15^ Phenotypic heterogeneity, driven by isogenic subpopulations that are not detected by standard, bulk population susceptibility tests widely used in the clinic, has been invoked as one potential explanation for this discrepancy.^16–18^

Non-genetic mechanisms that contribute to heterogeneity in drug response across the tree of life include tolerance, persistence, and heteroresistance.^7,19–26^ Although they are driven by diverse mechanisms, such states are typically transient and cannot be stably maintained in the absence of selective pressure.^7,9,24,25,27,28^ We previously reported a prion-based epigenetic mechanism in budding yeast that also gives rise to phenotypic heterogeneity.^8^ Unlike these other non-genetic mechanisms of adaptation, the acquired phenotypes are stably propagated through both mitotic and meiotic divisions without altering the genome. Yet how these mechanisms couple environmental cues to the production and propagation of phenotypic diversity, thereby promoting resilience and survival, is poorly understood. It is also unclear whether these mechanisms operate in other organisms, including evolutionarily distant fungal pathogens.

*Candida albicans*, a major contributor to the 3.8 million annual deaths caused by fungal infections^29^, serves as an excellent model for understanding how non-genetic sources of phenotypic variation influences adaptation. *C. albicans* thrives as both a commensal and a pathogen in humans and other mammalian hosts, exhibiting a high degree of plasticity in response to developmental cues and environmental stresses.^30–32^ Drug resistance in *C. albicans* is especially problematic for azoles, the antifungal class most widely used in the clinic, a substantial contributor to the ∼60% mortality rate associated with systemic infections. Azoles are also used extensively in agriculture, and persist in the environment at low levels due to their resistance to biodegradation.^33^ Although challenging to demonstrate experimentally, large-scale sequencing studies of *Aspergillus fumigatus* isolates from both patients and the local environment suggest that many infections likely acquired azole resistance before infecting the host.^34–36^ However, the molecular pathways and cellular processes that might enable the rapid emergence of drug resistance during exposure in this context have not been explored. Here, we combine microbial physiology, systems genetics, and cell biology to discover an inducible epigenetic state that integrates protein-based and chromatin-based mechanisms of inheritance to fuel the rapid evolution of drug resistance, with strong impacts on therapeutic efficacy both *in vitro* and in animal hosts.

## Results

### Robust and heritable adaptation to fluconazole in clonal populations

To investigate how *C. albicans* adapts to antifungals at the population level, we first exposed a sensitive laboratory strain (SN95) to fluconazole in a microbroth dilution-based minimum inhibitory concentration (MIC) assay.^37^ We inoculated ∼7,000 cells in rich liquid medium containing 2-fold serial dilution of the drug in 96-well plates. In wells without drug, *C. albicans* divided ∼12 times over 48 hours. As expected, exposure to fluconazole at the strain’s MIC (1.63 μM) impaired proliferation, with cells dividing ∼10 times over 48 hours (Fig 1A). We then asked how the descendants of pre-treated cells responded to a second round of drug exposure, repeating the MIC assay with fluconazole-treated inoculum. Cultures seeded with pre-exposed inoculum grew three times faster in medium containing fluconazole (1xMIC) than those inoculated of naïve cells (Fig 1A, Welch’s t test, ** p < 0.01). This phenotypic resistance was accompanied by a ∼2-fold increase in MIC (Fig S1A). That is, *C. albicans* can rapidly adapt to subtherapeutic doses of fluconazole at the population-level, and resistance is heritable over multiple divisions.

**Fig. 1.**
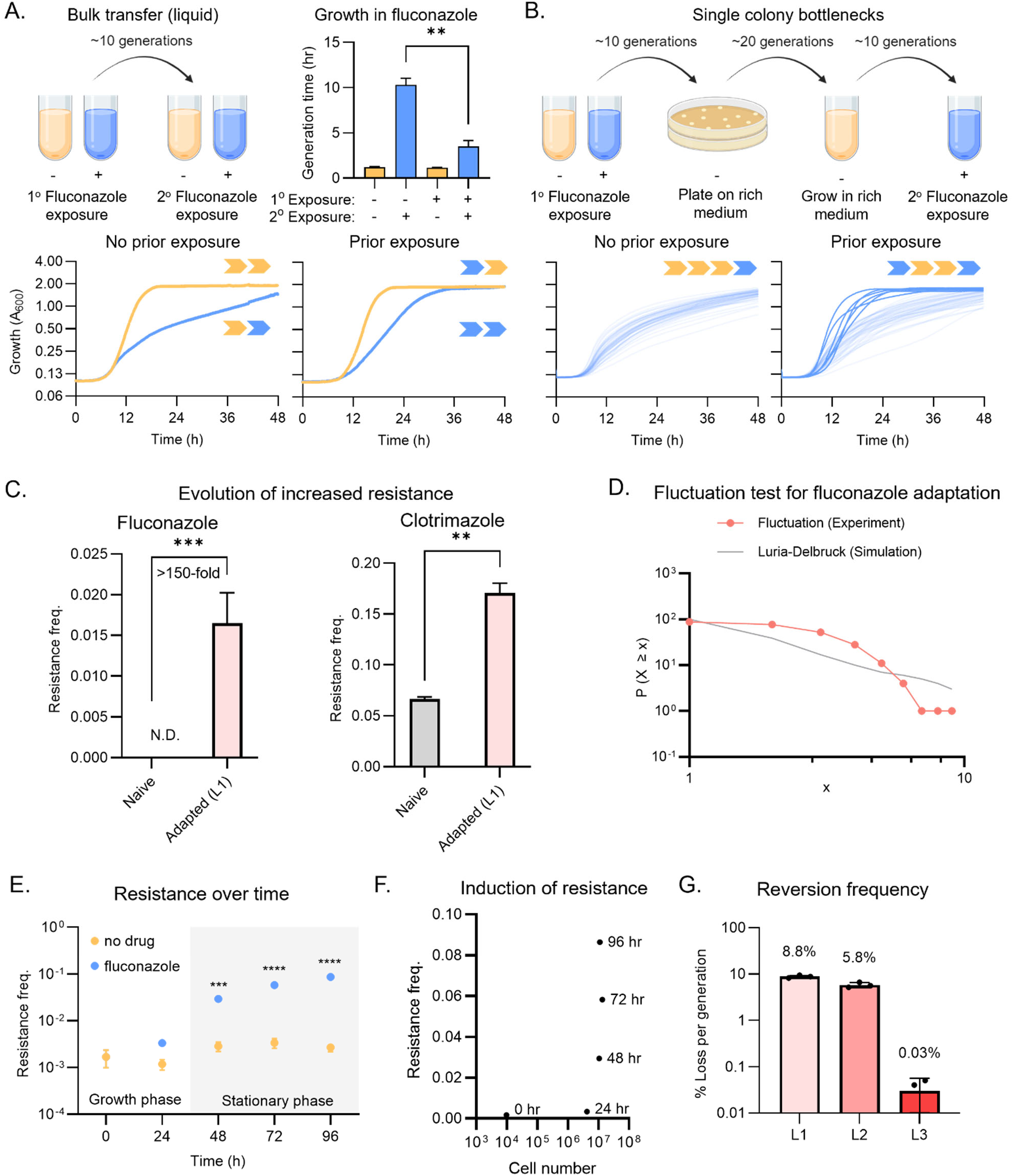
Subclinical doses of antifungal rapidly elicit heritable adaptation to drug. (A) Transient exposure to antifungal at near-MIC concentrations primes bulk adaptation to subsequent treatment in a laboratory *C. albicans* isolate. Growth of SN95 in YPD (yellow) and 1.63 μM fluconazole (blue) with no prior exposure to fluconazole (bottom left) and with prior exposure to fluconazole (bottom right). Yellow arrow indicates growth in YPD; blue arrow indicates growth in 1.63 μM fluconazole. The bar chart (top right) shows the generation time across two biological replicates. Error bars represent SD from two biological replicates. Welch’s t test, ** p < 0.01. (B) Memory of a prior exposure to antifungal promotes adaptation at high frequency in a clonal population. Growth of SN95 in 1.63 μM fluconazole. Light blue indicates no adaptation to fluconazole, dark blue indicates adaptive growth in fluconazole. Yellow arrow indicates growth in YPD; blue arrow indicates growth in 1.63 μM fluconazole. (C) Transient exposure to subinhibitory concentrations of fluconazole primes resistance to inhibitory doses of azoles. The bar chart shows the number of colony forming units on 6.53 μM fluconazole (left) and 742 nM clotrimazole (right). Gray represents naive SN95 and light pink represents the adapted lineage L1. Error bars represent SD from three technical replicates. Unpaired one-tailed t test, ** p < 0.01 and *** p < 0.001. (D) Adaptation to fluconazole did not arise spontaneously in untreated populations of *C. albicans* cells. The fluctuation test on 1.96 μM fluconazole across 100 technical replicates produces a distribution with a mean of 2.63 and a variance of 2.74. The solid gray line represents the predicted Luria-Delbruck distribution across 100 simulations, with a mean of 2.98 and a variance of 33.06. (E) Acquisition of the adaptive state depends on the duration of the drug exposure but not the number of divisions. Each data point represents an average of 10 technical replicates. Yellow indicates growth in YPD and blue indicates growth in 1.63 μM fluconazole. Two-way ANOVA, *** p < 0.001, **** p < 0.0001. (F) Phenotypic resistance continues to increase after the culture has saturated in fluconazole. Each data point represents an average of 10 technical replicates. Cultures were grown in 1.63 μM fluconazole. (G) The adaptive states are reversible. Light pink represents adapted lineage L1, salmon represents adapted lineage L2, and red represents adapted lineage L3. The bar chart shows the % loss per generation across the three technical replicates.

We next investigated whether phenotypic resistance could also arise in clinical isolates, examining a strain isolated from an HIV patient at the onset of oral candidiasis.^38^ Like in the laboratory strain, pre-exposure with fluconazole at MIC elicited bulk adaptation to future drug treatment (Figure S1B, top). We observed similar results in a bronchial isolate from a cystic fibrosis patient (Figure S1B, bottom). Thus, clonal *C. albicans* populations from distinct origins can adapt to subtherapeutic fluconazole concentrations via rapid acquisition of phenotypic resistance.

To better understand the rapid onset of these robust increases in fitness, we plated cells from four parallel untreated or fluconazole-treated cultures directly onto solid rich medium without drug. After two days of incubation, we picked 12 single colonies from each plate (96 in total) and used them to inoculate independent liquid cultures in fluconazole medium (1xMIC). All 48 lineages derived from the untreated cultures remained sensitive to drug after this single-colony bottleneck (Fig 1B, left). By contrast, six of the 48 lineages (12.5%) derived from the pre-treated cultures maintained resistance to fluconazole despite >30 generations of propagation without selection, during which the cells that had experienced the initiating stress were diluted by more than a billion-fold (Fig 1B, lower right). This observation suggests that the high frequency resistance to fluconazole is heritable through lineages over many generations, allowing the trait to persist within the population even after prolonged growth in drug-free medium.

We subsequently investigated how the adapted lineages responded to higher drug concentrations. To limit the confounding effects of increased basal resistance, we turned to an adapted lineage with the smallest increase in MIC (L1, < 2-fold relative to naïve; Fig S1C). Upon challenge with high doses of azoles, this lineage produced 150-fold more fluconazole resistant colonies (solid medium, 4xMIC), and 3-fold more colonies that are resistant to clotrimazole (solid medium, 16xMIC), a structurally distinct but mechanistically related antifungal (Fig 1C). Thus, transient exposure to subtherapeutic doses of fluconazole is associated with the rapid and frequent emergence of heritable phenotypic resistance in diverse *C. albicans* populations. Moreover, this adaptive state can significantly accelerate the evolution of resistance to higher concentration of multiple azole antifungals.

### Selection vs. induction during drug adaptation

This high frequency of adaptation could arise from selection on pre-existing variation or induction of *de novo* changes in response to drug. To estimate the influence of selection, we plated 94 independent cultures of naïve SN95 onto solid medium containing fluconazole at (1xMIC). Large colonies appeared at a frequency of ∼1.29 × 10^−3^ (Fig S1D). Cultures inoculated from these large colonies doubled 6.4-times faster in fluconazole than those inoculated from small colonies (Fig S1E and S1F, Welch’s t-test, p < 0.01), confirming that they were genuinely resistant to drug. Even under the conservative assumption that the adapted cells are entirely unaffected by drug and thus grow at rates equivalent to naïve cells in rich medium, a back-of-the-envelope calculation for an inoculum of 7,000 cells suggests that selection could have enriched resistant lineages to at most ∼0.03% of the population, ∼400-fold below the 12.5% we observed (Fig 1B).

We next performed a fluctuation test to examine the possibility that resistance was induced by fluconazole exposure.^39^ In such analyses, resistance that arises randomly during population growth yields highly variable numbers across multiple parallel experiments, including ‘jackpots’ with many resistant colonies. This scenario is described by a Luria-Delbrück distribution, where the variance greatly exceeds the mean.^39^ Alternatively, if resistance is induced in response to drug exposure, the number of resistant colonies across parallel experiments is comparable, conforming to a Poisson distribution where the variance and mean are similar.^39^ We grew 100 cultures of naïve SN95 in rich medium, inoculating each with <10 cells to avoid resistant founders, and after growing for 48 hours (to saturation; ∼20 divisions) plated them onto solid fluconazole medium (1xMIC). The resulting resistant colony counts conformed to a Poisson distribution with a similar mean and variance (Fig 1D; mean = 2.63 and variance = 2.74). Moreover, there were no jackpots, lending further support to the idea that high frequency adaptation to fluconazole arises in response to drug exposure.

Finally, we asked if such adaptation required cell proliferation, a hallmark of selection, or merely drug exposure, as might be expected for induction. We grew 10 independent liquid cultures with and without fluconazole for 96 hours, measuring both viable cell numbers and the frequency of resistance daily. By 36 hours, cell densities in both untreated and treated cultures had saturated. Untreated control cultures showed no difference in fluconazole resistance frequency over the remaining 60 hours in the experiment, during which no further proliferation was detected (Fig 1E). However, in drug-treated cultures, the resistance frequency continued to increase substantially (Fig 1F), depending on the duration of drug exposure rather than the number of cell divisions. Collectively, these data establish that induction by drug is a critical mechanism of phenotypic resistance to fluconazole in *C. albicans*.

### Phenotypic resistance is reversible

Given the rates of mutagenesis in *C. albicans* (10^−10^/bp/generation)^40^, genetic mutations could only explain the frequency of adaptation we observed under the implausible assumption that 15% of all base substitutions in the genome could confer dominant resistance to drug (Fig. S1G). This led us to suspect that resistance was not driven by genetic mutations. Indeed, when we used whole genome sequencing to examine L1 along with two other adapted lineages (L2 and L3), we observed no shared genetic changes, nor any variation in known drug resistance genes, even in individual lineages (Table S4).

The pronounced genome plasticity of *C. albicans* enables the frequent generation of aneuploidies that can confer stress adaptation and revert in the absence of selection. Moreover, long-term exposure to therapeutic doses of fluconazole has been associated with segmental and whole chromosome aneuploidies in laboratory selection in this organism.^41,42^ In one study, about half of resistant strains isolated from patients were aneuploid.^41–43^ Of the three adapted lineages we examined in detail, one was euploid and two harbored aneuploidies (Table S4), consistent with prior work.^41,42,44,45^ Although in other contexts, specific aneuploidies have been associated with adaptive drug responses, multiple lines of evidence suggest that this mechanism is not the primary cause of phenotypic resistance in our experiments: 1) We did not detect aneuploidy in all adapted isolates; 2) Phenotypic resistance does not require cell division but aneuploidy does;^46,47^ 3) Most aneuploidies confer a fitness cost in the absence of drug,^48^ yet we observed no correlation between increased phenotypic resistance and growth rate across 60 resistant lineages tested (Fig S1H); 4) In independently constructed strains with defined aneuploidies,^49^ the aneuploidy we observed most frequently did not confer resistance to fluconazole (trisomy Chr3 ABB, Fig S1I). The other aneuploidy that we observed infrequently (trisomy Chr5 ABB) has not been generated independently and has not emerged in previous studies examining the evolution of fluconazole resistance in this organism.^41,42,50^ Thus, although aneuploidy could, in principle, explain some properties associated with phenotypic resistance, it does not appear to be a dominant contributor to the phenotype in the lineages we examined.

The lack of genetic changes associated with the resistance phenotype, in combination with its high frequency and inducibility, pointed toward to epigenetic mechanisms, which can be heritable but typically revert more frequently than mutations.^51^ To examine phenotypic reversion, we compared clonal isolates of the adapted lineages to a resistant mutant control by streaking them for single colonies on solid fluconazole medium. Most progeny from each adapted lineage formed large colonies (Fig S1J); however, the streaks frequently contained a few small colonies, indicative of phenotypic reversion. By contrast, we never observed small colonies when streaking a fluconazole-resistant mutant (heterozygous hyperactive *UPC2* allele^52^, Fig S1J).

To quantify reversion frequency, we performed serial passage of lineages L1-L3 in liquid rich medium, measuring the frequency of resistance every ∼10 generations by plating onto solid fluconazole medium. Reversion frequencies ranged from 0.01% to 10% per generation for different adapted lineages. However, they were extremely reproducible in progeny from the same lineage (Fig 1G); for example, resistance was lost at 6.63%, 5.59%, and 5.18% per generation for three biological replicates of L2 and 0%, 0.04%, and 0.05% for L3. Together, these results suggest that the adaptive state acquired is not a simple binary switch; instead, it can exist as multiple stable variants that spans a spectrum in both resistance level and mitotic stability.

Phenotypic resistance to fluconazole thus combines environmental plasticity with the robust heritability classically associated with genetic changes. Yet the switch in cellular phenotype is reversible, a hallmark of epigenetic mechanisms. Multiple lines of evidence distinguish this behavior from tolerance, heteroresistance, and other mechanisms previously reported to increase subpopulation resistance in this organism.^25,53^ Thus, we adopt the term ‘para-resistance,’^54^ to describe this previously uncharacterized and highly unusual form of drug resistance.

### Multidrug resistance profile

We next investigated the diversity of the para-resistance phenotype, developing a colorimetric assay that allows rapid quantification of drug resistant colonies. Fluconazole permeabilizes the plasma membrane by targeting a critical step in ergosterol biosynthesis.^55^ We found that this was associated with selective accumulation of phloxine B dye, enabling a dual parameter Permeability-based Drug Resistance (PDR) assay that distinguishes between sensitive and resistant colonies by both size and color. Sensitive lineages produced uniformly small and dark colonies on PDR plates, whereas resistant mutants produced uniformly large and light colonies, with colors ranging from rose to white (Fig 2A, top row). By contrast, para-resistant linages produced large light colonies, as well as occasional small dark colonies, consistent with their capacity to revert to a susceptible state (Fig 2A, bottom row). Cultures inoculated with large light colonies (e.g. L3) consistently exhibited higher MICs than those inoculated with small dark colonies (e.g. naïve), establishing the assay’s capacity to discriminate between resistant and sensitive lineages (Fig 2A and S1C).

**Figure 2.**
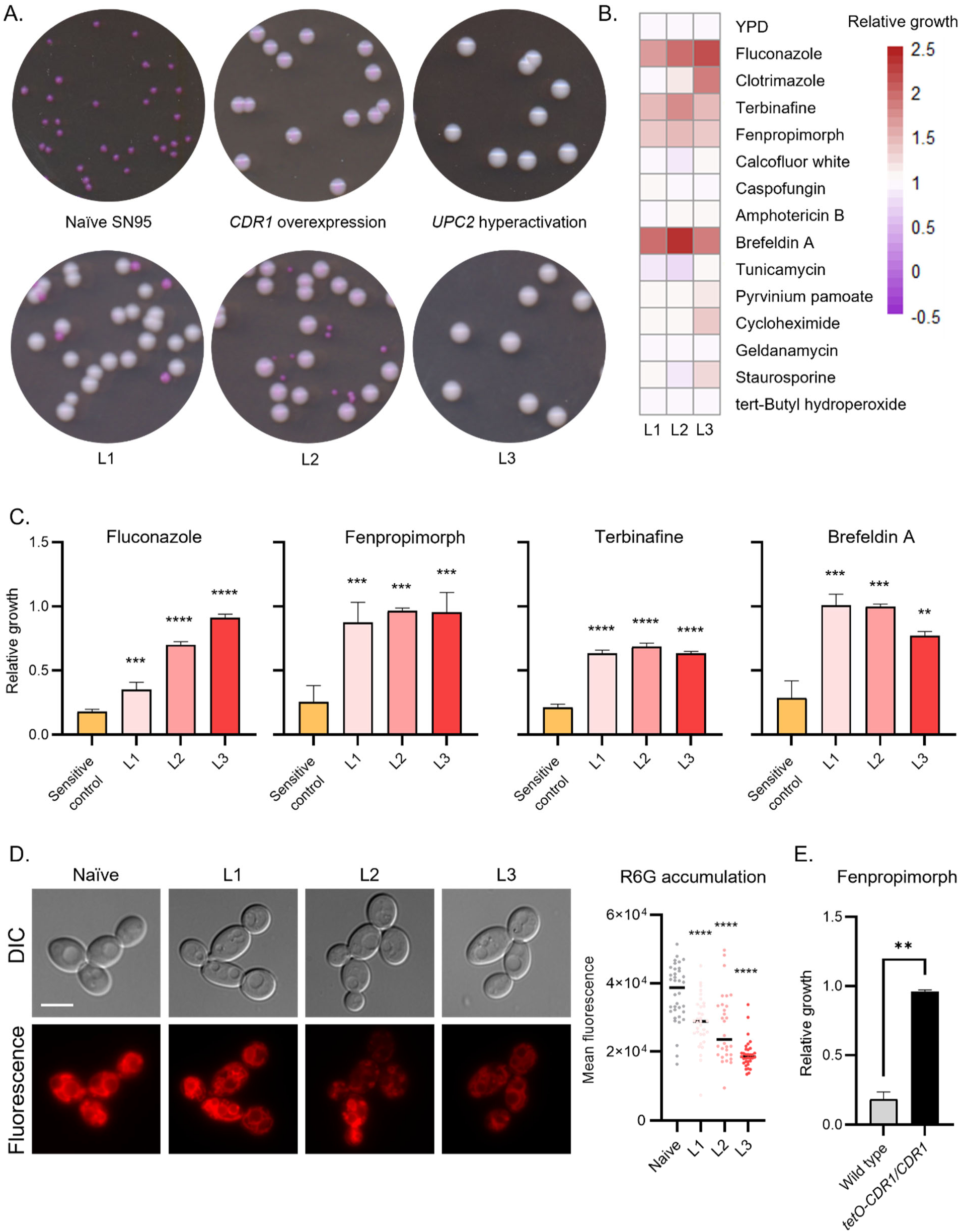
Para-resistant isolates are multi-drug resistant. (A) Phloxine B accumulation in the presence of fluconazole readily distinguishes sensitive from resistant colonies. Cells were plated on YPD plates supplemented with 1.96 μM fluconazole and 6.03 μM phloxine B. Plates were imaged after incubating for 48 h at 30°C. (B) Para-resistant lineages are resistant to multiple antifungal compounds. Phenotypic screen was conducted on three naïve and three para-resistant (L1-L3) isolates in four technical replicates across 14 conditions. Colony size on each condition was quantified using SGA tools. Growth of the sensitive and para-resistant isolates were normalized to the average growth of the naïve isolates. Data was quantitatively displayed as a heatmap (see color bar). (C) Para-resistant lineages share cross-resistance to fluconazole, terbinafine, fenpropimorph, brefeldin A. Growth at 1.63 μM fluconazole, 200 nM fenpropimorph, 6.86 μM terbinafine, and 57 μM brefeldin A were measured by absorbance at 600 nm after 24 h at 30°C, averaged across 2 biological replicates, and normalized to the no drug control. One-way ANOVA, ** p < 0.01, *** p < 0.001, **** p < 0.0001. (D) Less rhodamine 6-G accumulated in para-resistant isolates. Accumulation of rhodamine 6-G was visualized by fluorescence microscopy (left). Scale bar represents 5 μm. Mean fluorescence was quantified for at least 30 cells per sample by ImageJ (right). One-way ANOVA, **** p < 0.0001. (E) *CDR1* overexpression confers resistance to fenpropimorph. Growth at 200 nM fenpropimorph was measured by absorbance at 600 nm after 24 h at 30°C, averaged across 2 biological replicates, and normalized to the no drug control. Unpaired one-tailed t test, ** p < 0.01.

To obtain a comprehensive understanding of the phenotypic variability across para-resistant isolates, we screened three para-resistant and three naïve control lineages for cross-resistance to 14 compounds with diverse activities, including other classes of antifungals such as amphotericin B (a polyene) and caspofungin (an echinocandin) (Fig 2B). Beyond their shared fluconazole resistance, all para-resistant lineages showed robust cross-resistance to multiple compounds. These included the Erg2 and Erg24 inhibitor fenpropimorph (a morpholine), the Erg1 inhibitor terbinafine (a allylamine), and the ADP-ribosylation inhibitor brefeldin A (Fig 2C). These data demonstrate that para-resistance renders *C. albicans* recalcitrant to a wide range of antifungal therapeutics. That is, para-resistance is a multidrug resistant state.

Although these molecules are structurally unrelated, three of them (fluconazole, terbinafine, and brefeldin A) are known substrates of the Cdr1 drug efflux pump, which is frequently overexpressed in azole-resistant *C. albicans* isolated from human patients.^56,57^ To determine whether changes in Cdr1 efflux activity are linked to para-resistance, we measured the intracellular accumulation of rhodamine-6G, a non-toxic fluorescent Cdr1 substrate,^58^ observing reduced fluorescence signal in all three lineages (ranging from 1.3-fold to 1.9-fold relative to naïve controls; Fig. 2D, One-way ANOVA, p < 0.0001), consistent with increased Cdr1 activity. *CDR1* overexpression also conferred strong fenpropimorph resistance, hinting that this mechanism may indeed explain the multidrug resistance of para-resistant lineages (Fig 2E). Other examples of cross-protection were evident in some but not all para-resistant lineages, including adaptation to clotrimazole, cycloheximide, staurosporine, and pyrvinium pamoate (Fig 2B, z-score > 3). Given that cycloheximide is also a known Cdr1 substrate, these differences in sensitivity could arise from quantitative differences in Cdr1-dependent efflux, or they could be conferred by Cdr1-independent mechanisms. In summary, our results establish that enhanced drug efflux is likely an important contributor to para-resistance and its capacity to reduce the efficacy of multiple antifungal drugs. Moreover, the clonal variation among para-resistant lineage across multiple parameters demonstrates the capacity of para-resistance to manifest as multiple stable variants with distinct multidrug resistant profiles.

### Chromatin-based and protein-based drivers

Phenotypic heterogeneity is classically understood as a property of epigenetic phenomena, ranging from position effect variegation to prion polymorphs.^59,60^ To find out if either of these epigenetic mechanisms could explain the inducibility, reversible heritability, and phenotypic diversification of para-resistant lineages, we asked whether small molecules that perturb chromatin modification and prion inheritance would impact induction of para-resistance. Two of the molecules that we tested blocked the induction of para-resistance: guanidinium hydrochloride, which blocks the propagation of fungal prions, and 1,6-hexanediol, which has been reported to dissolve liquid-liquid phase-separated condensates (52-fold and 62-fold; Fig 3A, One-way ANOVA, p < 0.0001). These results implicate prion-like protein assembly mechanisms in para-resistance.

**Figure 3.**
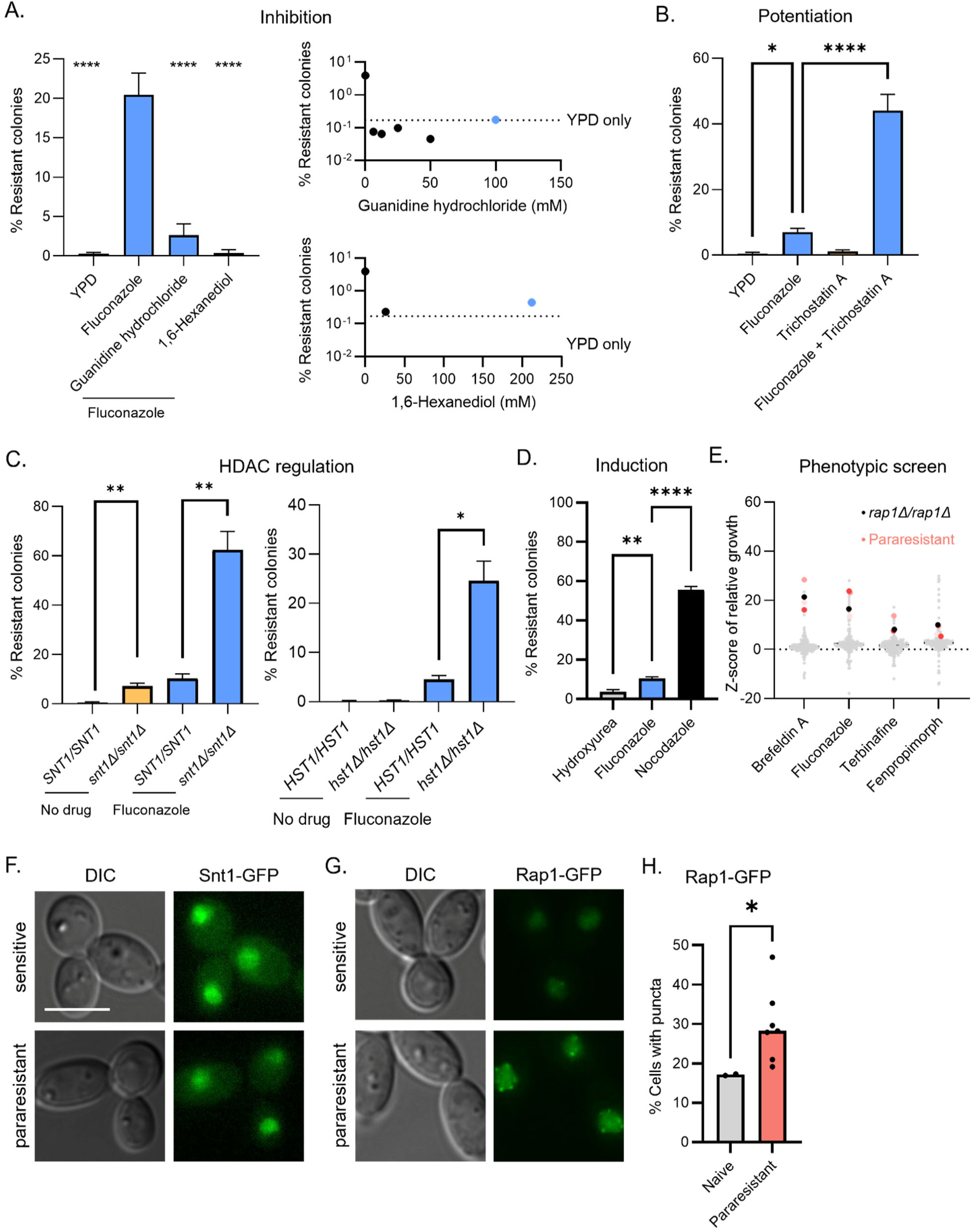
Chemical and genetic modulators of para-resistance. (A) Guanidine hydrochloride and 1,6-hexanediol block fluconazole-induced phenotypic resistance. (Left) Wild-type SN95 was transiently exposed to 1.63 μM fluconazole alone or in combination with 100 mM guanidine hydrochloride or 212 μM 1,6-hexanediol were plated on PDR plates. The bar plot shows the average of 5 technical replicates. One-way ANOVA, **** p < 0.0001. Guanidine hydrochloride (top right) and 1,6-hexanediol (bottom right) block the induction of para-resistance in a concentration-independent manner. Blue indicates the concentrations used in the bar chart (left). (B) Trichostatin A potentiates fluconazole-induced phenotypic resistance. Wild-type SN95 transiently exposed to 1.63 μM fluconazole, 20 μM trichostatin A, or both were plated on PDR plates. The bar plot shows the average of 3 technical replicates. One-way ANOVA, * p < 0.05, **** p < 0.0001. (C) Genetic perturbation of the Set3C histone deacetylase complex potentiates fluconazole-induced phenotypic resistance. Wild-type SN95 strain was transiently exposed to 1.63 μM fluconazole, *snt1Δ/snt1Δ* mutant (left) was transiently exposed to 0.98 μM fluconazole, and *hst1Δ/hst1Δ* mutant was transiently exposed to 1.63 μM fluconazole (Right). Cells were plated on PDR plates. The bar plot shows the average of 3 technical replicates. One-way ANOVA, * p < 0.05, **** p < 0.0001. (D) Nocodazole induces phenotypic resistance to fluconazole at high frequency. Wild-type SN95 transiently exposed to 1.63 μM fluconazole, 25 μM nocodazole, or 25 mM hydroxyurea were plated on PDR plates. The bar plot shows the average of 3 technical replicates. One-way ANOVA, ** p < 0.01, **** p < 0.0001. (E) Deletion of *RAP1* phenocopies the multidrug resistance phenotypes associated with para-resistant isolates. The transcription factor deletion mutants were grown on 1.96 μM fluconazole, 35.7 μM brefeldin A, 34.3 μM terbinafine, and 200 nM fenpropimorph. Colony sizes were quantified using SGA tools and normalized to Gray indicates transcription factor deletion mutant, black star indicates *rap1Δ/rap1Δ* mutant, light pink indicates para-resistant lineage L1, salmon indicates para-resistant lineage L2, and red indicates para-resistant lineage L3. (F) Snt1 localizes to the nucleus in naïve and para-resistant isolates. GFP-tagged Snt1 was detected by fluorescence microscopy. Scale bar represent 5 μm. (G) Rap1 localizes to the nucleus in naïve and para-resistant isolates. GFP-tagged Rap1 was detected by fluorescence microscopy. (H) Para-resistant isolates show an increased propensity for the formation of nuclear Rap1 puncta. At least 100 cells in 5 fields of view were visually screened and manually counted for nuclear puncta. Welch’s t test, * p < 0.05.

A third molecule, the histone deacetylase inhibitor trichostatin A, amplified the capacity of fluconazole to induce para-resistance (5.3-fold; Fig 3B, One-way ANOVA, p<0.0001). In *C. albicans*, trichostatin A inhibits the Set3/Hos2 histone deacetylase complex, composed of four core subunits (Set3, Hos2, Snt1, and Sif2) and three peripheral subunits (Hst1, Cpr1, and Hos4).^61,62^ Mutants eliminating a critical scaffold (*snt1Δ/snt1Δ*) and the complex’s deacetylase activity (*hst1Δ/hst1Δ*) acquired para-resistance more frequently when induced with fluconazole (6.1-fold and 5.8-fold; Fig 3C), suggesting that the loss of Set3C function promotes para-resistance.

Why does para-resistance show strong signatures of both prion-based and chromatin-based properties? In the model yeast *Saccharomyces cerevisiae*, the Set3C core subunit Snt1 can adopt a self-templating protein conformation, which can be triggered by phosphorylation in response to prolonged G2/M.^8^ This [*ESI^+^*] prion also confers resistance to fluconazole. Therefore, we tested whether prolonged G2/M arrest might induce para-resistance in *C. albicans* despite the 300 million years of evolutionary divergence that separates these two species.^63^ Indeed, the same nocodazole-induced G2/M arrest that triggered [*ESI^+^*] increased the frequency of para-resistance, and did so even more strongly than fluconazole (56% [n = 530] vs 10% [n =1213]; Fig 3D, One-way ANOVA, p < 0.0001). Induced variants spanned the full spectrum of phenotypes associated with spontaneous and drug-induced para-resistance (Fig S2A). By contrast, treatment with hydroxyurea, which elicits an S-phase arrest,^64^ did not significantly affect the induction of para-resistance, establishing that the effect of nocodazole was not due to a generic cell cycle slowdown (Fig 3D).

In [*ESI^+^*] cells, reorganization of Set3C is associated with eviction of Rap1 (Repressor-activator protein) from sub-telomeric HAST (Hda1-affected sub-telomeric) domains, which are normally silent in *S. cerevisiae*.^8,65^ Conditional depletion of Rap1 from the chromatin phenocopies many of the [*ESI^+^*] phenotypes.^8^ Two lines of evidence suggest that loss of Rap1 function also plays a critical role in para-resistance. First, when we screened 165 transcription factor deletion mutants,^66^ *rap1Δ/rap1Δ* was the only mutant that conferred cross-protection to fluconazole, brefeldin A, terbinafine, and fenpropimorph, phenocopying the three para-resistant lineages described above (Fig 3E; within 1.5 SD of L1-L3). Second, just as we observed with nocodazole, *rap1Δ/rap1Δ* colonies were resistant to fluconazole, displaying a range of sizes and colors that encompassed the spectrum observed across diverse para-resistant lineages (Fig S2B). Collectively, our findings suggest that conserved molecular mechanisms underlying [*ESI^+^*] are related to those involved in para-resistance, and by analogy, inheritance of the phenotype likely integrates prion-based mechanism to stabilize an epigenetic memory driven by chromatin modification.

### Cell biological consequences

Both Rap1 and Snt1 are DNA-binding proteins. Therefore, we tested whether changes in their subcellular localization are associated with the acquisition of para-resistance. We fused GFP to the C-termini of Rap1 and Snt1 to compare their distributions in naïve and para-resistant isolates using fluorescence microscopy. Snt1-GFP localized to the nucleus (Fig 3F) and its fluorescence increased significantly in naïve cells following fluconazole exposure (Fig S2C, Welch’s t test, p< 0.001). However, the nuclear-to-cytoplasmic ratio (often used as a proxy for activity) of Snt1-GFP was reduced, rather than elevated, in most para-resistant isolates (Fig S2D, Welch’s t test, p < 0.05; similar mean fluorescence per cell).

Next, we studied Rap1-GFP in seven independent para-resistant lineages. In naïve lineages, Rap1-GFP was mostly diffuse, with occasional cells showing small numbers of weak puncta (Fig 3G). In contrast, most para-resistant lineages exhibited both higher proportion of cells with Rap1-GFP puncta and brighter, more numerous puncta in those cells that contained them (5 of 7, Welch’s t test, p < 0.05; Fig 3G-H). Sensitive revertants derived from these lineages lost Rap1-GFP foci and resembled naïve cells (Fig S2E). These observations indicate that a heritable change in Rap1 function and localization are common features among para-resistant lineages.

To investigate whether the Rap1-GFP puncta were associated with conformational changes in Rap1, we performed limited proteolysis^67^ on Rap1-GFP in lysates from a naïve control and two para-resistant isolates exhibiting medium or high levels of Rap1-GFP puncta (Fig S2F). Following proteinase K treatment, a stable ∼55 kDa fragment accumulated as the full-length Rap1-GFP was degraded. Interestingly, this proteolytic processing was more efficient in para-resistant cells (Fig S2G), perhaps reflecting a structural rearrangement of Rap1’s untagged, N-terminal disordered region. Together, our results reveal that stable propagation of altered Rap1 conformations is associated with para-resistance. Combined with prior lines of genetic and chemical evidence, our cell biological and biochemical observations indicate that para-resistance involves a structural and functional remodeling of chromatin silencing factors, including Rap1, generating a multi-layered regulatory network that enables rapid and stable adaptation to antifungal stress in *C. albicans*.

### The para-resistant transcriptional program

Since histone acetylation is often associated with increased gene expression, we investigated the transcriptional signatures of para-resistance using mRNA-sequencing on three naïve and three para-resistant populations in the presence and absence of fluconazole. The transcriptomes of the para-resistant isolates overlapped and differed from those of untreated cells and of pretreated but non-adapted lineages, even in the absence of fluconazole (704 differentially expressed genes with at least 2-fold change in L1, 1,298 in L2, 630 shared; Fisher’s exact test, OD = 55.9241, p<0.0001; Fig 4A, Fig S3A and Table S5). The transcriptome of the third lineage differed less markedly (21 differentially expressed genes in L3; Fig S3A and Table S5). This lineage had no detectable SNPs, confirming its non-genetic origin, likely involves a different underlying mechanism (Table S4 and Fig 2B).

**Figure 4.**
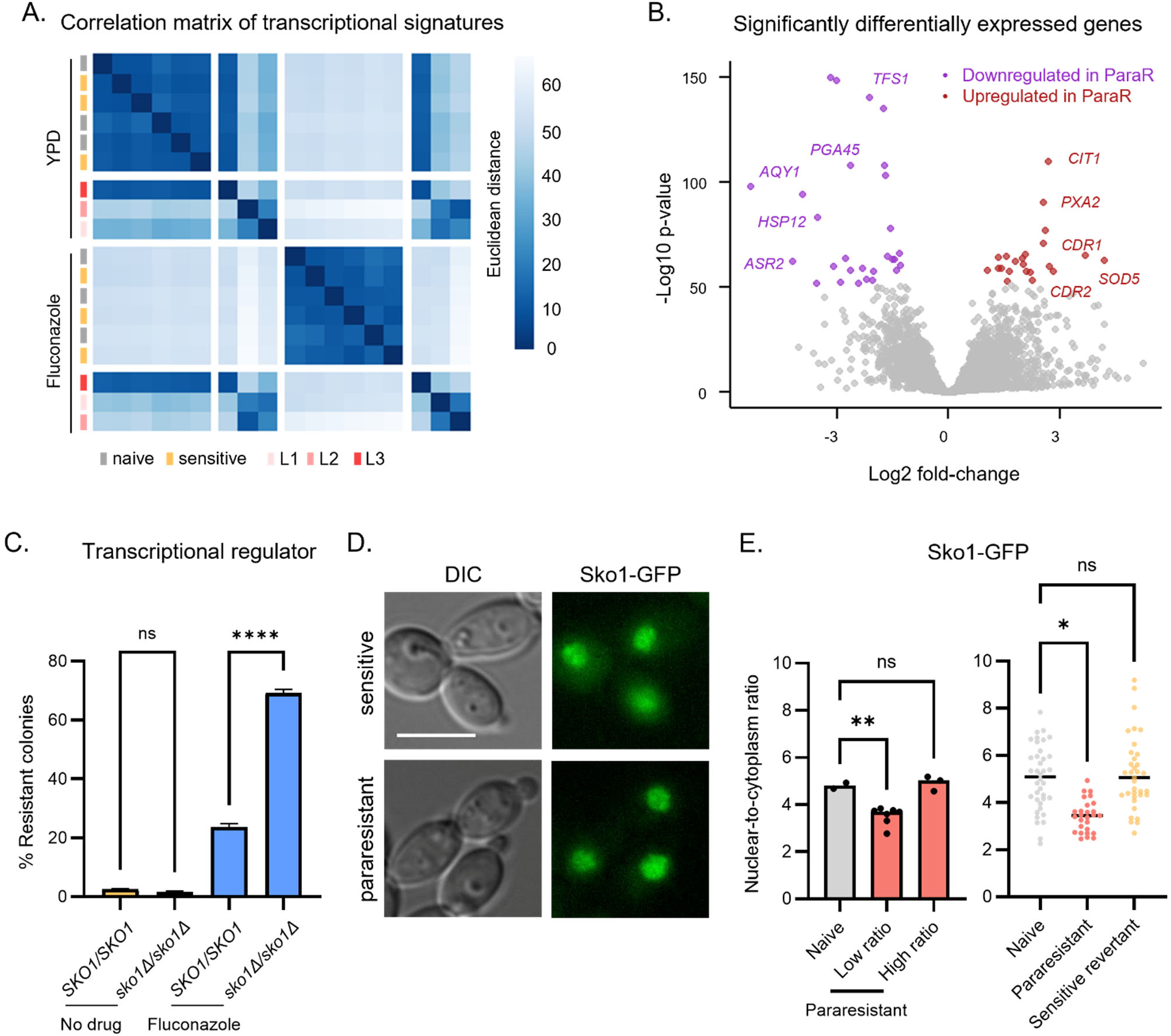
Transcriptional regulation of para-resistance. (A) Correlation matrix of the gene expression signatures associated para-resistance and fluconazole treatment. Heat map highlighting the similarities and differences in transcriptional signature among naïve (N1-N3), sensitive (S1-S3), and para-resistant isolates (L1-L3). (B) Volcano plot of genes significantly differentially expressed between para-resistant and sensitive isolates. The top 50 significantly differentially expressed genes are highlighted in red (upregulated in naïve) and purple (downregulated in naïve). (C) Deletion of *SKO1* potentiates fluconazole-induced phenotypic resistance. Wild-type SN95 and *sko1Δ/sko1Δ* strains were transiently exposed to 1.63 μM fluconazole. Cells were plated on PDR plates. The bar plot shows the average of 3 technical replicates. One-way ANOVA, **** p < 0.0001. (D) Sko1 localizes to the nucleus in naïve and para-resistant isolates. GFP-tagged Sko1 was detected by fluorescence microscopy. Scale bar represent 5 μm. (E) Sko1 changes in nuclear distribution in some of the para-resistant isolates. Fluorescence signal in the nucleus and cytoplasm was quantified using ImageJ for naïve (2) and para-resistant (7/10) isolates and plotted as a ratio (left). Welch’s t test, ** p < 0.01. The nuclear-to-cytoplasmic ratio was calculated for at least 27 cells per sample by ImageJ (right). One-way ANOVA, * p < 0.05, ** p < 0.01.

Whereas exposure to fluconazole elicited strong transcriptional changes in both naïve and non-adapted controls, as expected from prior studies of *C. albicans* responses to fluconazole^68^, the transcriptome of the para-resistant linages were largely unaffected by drug (Fig 4A and Fig S3B). The para-resistant lineages also did not show any transcriptional evidence of pre-adaptation to drug or constitutive activation of stress responses.^69^ In fact, stress-responsive heat shock genes such as *ASR1* and *HSP12* were significantly repressed (Fig 4B). Moreover, only two of the fifty most differentially expressed genes, which encode multidrug transporters (*CDR1* and *CDR2*)^70^ are known to impact fluconazole resistance (Fig 4B). These observations establish that para-resistance can arise from drug refractory epigenetic states driven by multiple mechanisms, and often involves global transcriptional changes. None appears to be driven by canonical stress response pathways.

Next, we searched for regulators of the shared transcriptional response in L1 and L2. We queried the compendium of regulatory associations between transcription factors and target genes in pathogenic fungi (PathoYeastract)^71^, using sliding windows encompassing the top 50 – 500 most differentially expressed genes in the two lineages. This analysis identified five putative transcriptional regulators of: Ace2, Cta8, Sko1, Efg1, and Mac1 (Table S6). Sko1-regulated genes were most strongly overrepresented (>2.5-fold enrichment) among those differentially expressed in para-resistant vs naïve transcriptomes.

We assessed how deleting each of these transcription factors affected growth in fluconazole (apart from *CTA8*, which is essential). Deletion of two factors (*EFG1* and *ACE2*) increased colony size by >1.5-fold (Fig S3C). However, neither produced the spectrum of colony color and size characteristic of para-resistance, nor did they phenocopy the multidrug resistance profile shared among L1-L3. Cells harboring *MAC1* deletion produced small colonies, but their normalized susceptibility to fluconazole was equivalent to a naïve control. Thus, para-resistance does not appear to arise from simple gains or loss of these functions. Interestingly, *sko1Δ/sko1Δ* mutants did not affect growth on fluconazole (Fig S3C), but the frequency of fluconazole-induced para-resistance increased substantially in this mutant (39.9-fold; Fig 4C, One-way ANOVA, p < 0.0001), linking its function to the acquisition of para-resistance.

Sko1 contains extensive intrinsically disordered regions (IDRs). Biases within these sequences resemble those in some self-templating prion proteins (Fig S3D). We therefore examined Sko1-GFP levels and localization, observing predominantly nuclear signal at similar levels in naïve and para-resistant cells, consistent with its function a transcription factor (Fig 4D and Fig S3E). However, the ratio of nuclear-to-cytoplasmic signal was significantly lower in most para-resistant lineages (7 of 10, Fig 4E, left). Interestingly, nuclear-to-cytoplasmic Sko1-GFP ratios returned to naïve levels in a sensitive revertant derived from a para-resistant isolate that initially had low ratios (Fig 4E, right). These results suggest that changes in the nuclear distribution of Sko1, consistent with altered transcriptional function, appear to be associated with a majority of the para-resistant states.

We also searched for transcription factors whose binding sites are enriched among the differentially regulated genes using Analysis of Motif Enrichment (AME) with JASPAR CORE pan-fungal motif database. Analysis of the promoters of the 50 most differentially expressed genes in para-resistant lineages revealed significant motif enrichment for 11 DNA-binding transcription factors present in *C. albicans*, including Rap1 (Table S7). Interestingly, the strongest multicopy suppressor of toxicity associated with *RAP1* overexpression in *S. cerevisiae* is Sko1.^72^

Sko1 functions downstream of the High-Osmolarity Glycerol (HOG) pathway, and is phosphorylated by the Mitogen-Activated Protein Kinase (MAPK) Hog1 in response to osmotic stress.^73^ Strikingly, in the *hog1Δ/hog1Δ* mutant cells, para-resistance induction frequency was reduced 3-fold (Fig S3F), suggesting that this stress response circuitry may prime the acquisition of para-resistance when conditions are unfavorable.

### Emergence of para-resistance *in vivo*

In the clinic, drug susceptibility is typically measured as the MIC, the lowest drug concentration that inhibits visible microbial growth.^37^ Resistance, often indicated by an increase in MIC, is commonly attributed to genetic mutations.^74^ However, previous studies have identified transient fluconazole resistance in a series of clinical *C. albicans* isolates associated with treatment failure, suggesting that heritable and reversible resistance may be more prevalent in the clinic than previously recognized.^75–77^ To ask if para-resistance could be hidden in plain sight among clinical isolates, we obtained *C. albicans* isolates from 9 cystic fibrosis patient samples at the Stanford Hospital (5 sputum, 2 body fluid, 1 urine, and 1 bronchoalveolar lavage). Five of these samples came from fluconazole-treated patients, but all had been classified as susceptible based on MIC (Table S8).

When we plated these isolates on PDR plates, we discovered that bulk MIC susceptibility measurements had hidden substantial variation in drug response. The frequency of spontaneous resistance to fluconazole was ∼10-fold higher in two sputum isolates (CaCiS-1, 2.8%; CaCiS-7, 3.1%) than in SN95, a naïve laboratory strain (0.32%). Interestingly, CaCiS-1 was isolated from a patient that had been treated with fluconazole, whereas CaCiS-7 was isolated from a patient that had not been prescribed an antifungal (Table S8). To assess whether this phenotypic resistance resembled para-resistance, we plated CaCiS-1 and CaCiS-7 onto rich solid medium and picked 30 colonies from each isolate. We then re-suspended these colonies and plated them onto PDR plates. Despite the single-colony bottlenecks inherent to this procedure, the derived colonies gave rise to both large resistant colonies and small sensitive colonies on PDR plates (Fig 5A and Fig S4A, top). As we previously observed in laboratory strains, the large pink and white colonies grew better in fluconazole than the small pink colonies (Fig 5A and Fig S4A, bottom). CaCiS-7 derivatives gave rise to colonies that were homogenous in color and size (Fig 5B and Fig S4A); however, CaCiS-1 produced more heterogenous colonies, providing evidence for fluconazole resistance that is both heritable and reversible – a defining feature of para-resistance (Fig 5A).

**Figure 5.**
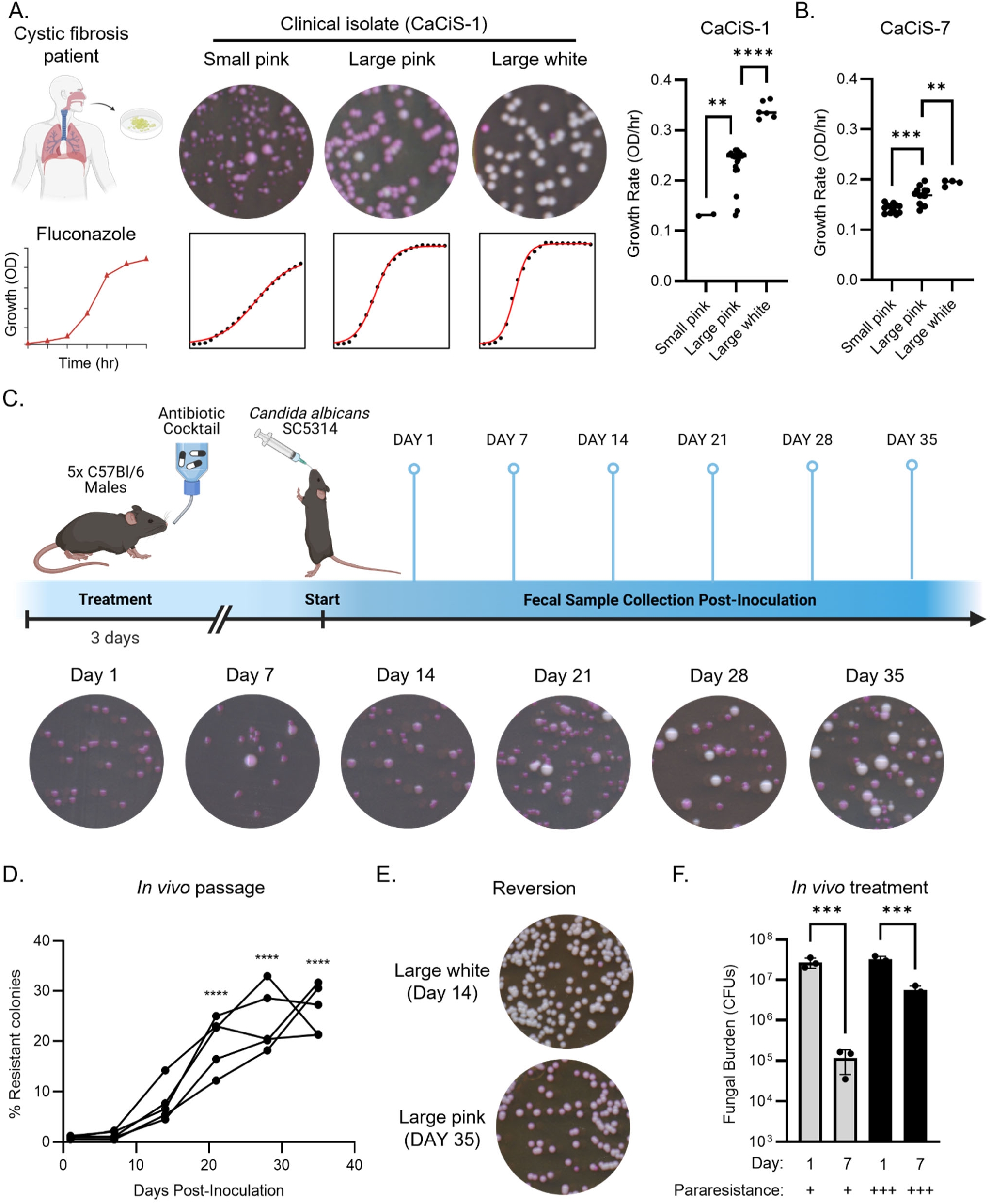
Para-resistance acquired *in vivo* diminishes antifungal therapeutic efficacy. (A) and (B) Para-resistance is detected in a clinical *C. albicans* isolate. CaCiS-1 (isolated from a cystic fibrosis patient) was plated on YPD plates. 30 single colony derivatives were plated on PDR plates. Growth in 3.62 μM fluconazole was measured over 96 h. The bar plots show the growth rate for each colony type derived from CaCiS-1 (A) and CaCiS-2 (B). One-way ANOVA, ** p < 0.01, *** p < 0.001, **** p < 0.0001. (C) Schematics for mouse gastrointestinal colonization with *C. albicans*. SC5314 was administered to 5 antibiotic-treated mice at 10^7^ cells. Mouse fecal samples were collected every 7 days and plated for single colonies on PDR plates. The plates were imaged after 48 h incubation at 30°C. (D) High frequency fluconazole resistance emerges with passage in the mouse gastrointestinal tract. Mouse fecal samples were collected every 7 days and plated for single colonies on PDR plates. The plates were imaged after 48 h incubation at 30°C. One-way ANOVA, compared to day 1 post-inoculation. * p < 0.05, ** p < 0.01. (E) Resistance acquired via *in vivo* passage is both heritable and reversible. Fecal samples from day 14 and day 35 post-inoculation were plated on YPD. Single colony derivatives were subsequently plated on PDR plates. The plates were imaged after 48 h incubation at 30°C. (F) Fluconazole is less effective in eliminating *C. albicans* cells that have acquired a high level of para-resistance. Mice were treated with 100 mg/kg/day. Fecal samples from day1 and day 7 post-treatment were plated on PDR plates. The plates were imaged after 48 h incubation at 30°C and colonies were counted manually.

To ascertain whether the phenotypic resistance in these strains arose from a mutational mechanism, we analyzed whole genome sequences for six CaCiS-1 derivatives (strains a-f) that form three classes (Table S9). Class I produced small dark colonies, class II produced large pink colonies, and class III produced large white colonies on the PDR plates, which corresponded to susceptible, moderate or strong resistance to fluconazole, respectively. No point mutations were evident in either class I or class III strains, despite their phenotypic differences (Table S9): strain a (class I) and strain f (class III) appear genetically identical, both carrying an extra copy of Chr3 B; however, strain a was more sensitive than strain f, which was resistant to fluconazole. Just in defined laboratory reconstruction, the Chr3 ABB trisomy is not sufficient to confer fluconazole resistance. By contrast, in strain d (class II), we identified one non-synonymous mutation in *TAC1*, which encodes a transcription factor that positively regulates *CDR1* and *CDR2* expression, and is known to affect responses to azole drugs^70^. This may explain why large pink colonies are resistant but cannot account for the high-frequency emergence of white colonies on PDR plates in this strain (Fig 5A).

Lastly, we evaluated the six CaCiS-1 derivatives for cross-resistance to other drugs. All six exhibited a range of multidrug resistance profiles (Fig S4B), leading us to suspect that para-resistance states often emerge in the clinic, are missed by current assays used to detect resistance, and may impede effective treatment.

### Gastrointestinal colonization

The detection of an apparently para-resistant strain from a patient who had not been treated with fluconazole (CaCiS-7) suggested that this phenomenon might emerge naturally in response to host-relevant stresses. *C. albicans* is a natural member of the human microbiota and a key player in immune priming.^78^ Sequencing at multiple body sites within the same individual suggests that a single strain can predominate.^79,80^

Given that *C. albicans* is the most frequently observed fungal species in the gastrointestinal tract,^81^ we used a well-established murine model of gastrointestinal colonization^82^ to monitor the emergence and dynamics of para-resistance. We orally inoculated 5 antibiotic-treated mice with the fluconazole-susceptible lab strain (SC5314) and subsequently collected fecal samples every 7 days (Fig 5C). We plated these samples onto PDR plates, quantifying colony number, size, and color (Fig 5C). Strikingly, the frequency of fluconazole resistance increased significantly during host passage (Fig 5D; Two-way ANOVA, p < 0.0001). After 35 days, 21% to 32% of the *C. albicans* cells recovered from each of the five mice were resistant to fluconazole (Fig 5D), establishing that host adaptation during commensalism is sufficient to trigger a drug-resistant state in substantial proportion of the population.

The phenotypic resistance that emerged *in vivo* shared several key properties with para-resistance acquired *in vitro*. For example, large colonies ranged in color, from different hues of pink to white, on the PDR plates and grew better on fluconazole (Fig 5B), with at least a 2-fold increase in MIC compared to the inoculum. Lineages propagated true to type (i.e. large white colonies gives rise to large white colonies), and were capable of reverting to produce small dark colonies (Fig 5E). Thus, like para-resistance induced *in vitro*, phenotypic resistance derived from *in vivo* passage is stable, yet reversible.

Lastly, we asked whether fluconazole resistance that arose spontaneously *in vivo* could confer resistance to therapeutic doses of fluconazole. We repeated the gastrointestinal colonization experiment with 12 mice, split into three cages. After 42 days, the frequency of para-resistance was especially high in one of the cages (71.7% ± 11.5%), and lower in the remaining two cages (2.5% ± 0.6% and 2.4% ± 0.3%), despite similar fungal burden (Fig S4D). We subsequently treated the mice with therapeutic doses of fluconazole (100 mg/kg/day). In mice with a low levels of para-resistance, we observed a 230-fold reduction in fungal burden after 7 days of treatment (Fig 5F, One-way ANOVA, p < 0.05). By contrast, the same drug treatment reduced fungal burden by just 6-fold in mice with high levels of para-resistance (Fig 5F, One-way ANVOA, p < 0.05). These data establish that passage through the gut of mammalian hosts can give rise to substantial para-resistant *C. albicans* that limit fluconazole therapeutic efficacy.

## Discussion

Across diverse biological systems and contexts, extensive phenotypic variation generated de novo and in response to drug-induced stress can facilitate the rapid appearance of strains that are able to grow in otherwise inhibitory drug concentrations.^83^ Research into the origins and physiological impact of such variation has primarily focused on stress response networks and genome diversification. Stress responses provide a transient yet powerful protective mechanism, creating a critical window of opportunity during which new variation can improve chances of survival.^84,85^ Likewise, stress can be coupled to the induction of mutagenesis^86^ to increase the number of variants on which natural selection can act on.^87^ Recent studies suggest that epigenetic mechanisms might also support this broad spectrum of adaptive potential.

Here, using *C. albicans*, a major human fungal pathogen, we found that transient exposure to fluconazole significantly increases adaptive variation that is not accompanied by mutations. This unusually high frequency of adaptation (∼1 in 10) also cannot be explained solely by selection on pre-existing variation. Rather, our data suggest that stable but reversible adaptive variation is created *de novo*, in response to drug. This dynamic epigenetic state strikes a balance between the stability required to maintain successful traits and the flexibility needed for rapid adaptation to environmental changes. Thus, heritable and reversible variation has the potential to drive the rapid evolution of adaptive traits.

From microbes to human cancers, a continuum of non-genetic mechanisms including tolerance, persistence, and heteroresistance, influence drug adaptation.^7,23,25,88^ Like para-resistance, these states do not rely on the acquisition of mutations and can thus arise more frequently than mutations, thereby enabling a disproportionate impact on survival. Yet para-resistance differs from each of these states in key physiological features and underlying mechanisms. Intrinsic antifungal tolerance is an inherent property of a strain, with tolerant and non-tolerant cells producing similar ratios of each cell types.^25,89^ Acquired antifungal tolerance often results from aneuploidy, copy number variation, and/or loss of heterozygosity.^42,90^ Antifungal heteroresistance occurs when a rare subpopulation of cells transiently displays an MIC that is 8- to 10-fold higher than the parent population.^53,89^ The progeny cells revert to a drug-susceptible state when grown without selective pressure.^91^ By contrast, para-resistance is far more stable than these other drug adapted states, propagating faithfully over dozens to hundreds of mitotic divisions. Moreover, para-resistance is induced by the selective pressure against which it provides resistance, and once acquired, enables adaptation to much higher doses of drug. We posit that this unusual constellation of properties arises from an interplay between protein-based and chromatin-based epigenetic mechanisms, enabling cells to integrate their decision to switch with signals from deeply conserved stress-sensing circuitry.

Although epigenetic states are often established in response to stress, the mechanisms by which cells retain a ‘memory’ of prior stress that facilitates adaptation to future challenges often remain elusive. In the case of para-resistance, we found that several factors are involved: the stress-activated MAPK Hog1 positively regulates para-resistance, whereas its downstream transcription factor, Sko1, negatively regulates it. Furthermore, para-resistance and its strength are associated with increasing assembly of Rap1. Other observations in *S. cerevisiae* suggest broad conservation in relationship between Sko1 and chromatin activation/suppression. Notably, Sko1 is the strongest multicopy suppressor of toxicity associated with *RAP1* overexpression.^72^ Moreover, *RAP1* deletion confers resistance to fluconazole, brefeldin A, terbinafine, and fenpropimorph, which mirrors the multidrug resistance phenotype we observed in multiple para-resistant isolates induced *in vitro*. Thus, Rap1 is likely critical for the emergence of most para-resistant populations.

We previously made similar observations in *S. cerevisiae*, where reducing Rap1 function phenocopied [*ESI^+^*], a heritable and reversible prion state formed by conformational change in Snt1.^8^ Also consistent with findings in *S. cerevisiae*, chemical or genetic perturbation of the Set3C histone deacetylase function in *C. albicans* promotes the acquisition of the heritable and reversible para-resistance state. One working hypothesis is that cell-to-cell variation in Hog1 activation primes this switch by phosphorylating Sko1, which is a strong repressor of Rap1 in other organisms.^72^ Although additional mechanistic work is required to fully understand the diversity of the para-resistance phenotype, we propose that many of these processes may be fertile ground for therapeutic development.

Drug resistance mechanisms that evade standard surveillance confound the accurate prediction of treatment outcomes.^25,92^ Notably, isolates classified as susceptible in clinical drug susceptibility testing can nonetheless cause infections that are not cleared in patients despite treatment with inhibitory drug concentrations.^57,93^ Acquisition of pararesistance in laboratory and clinical strains increase MICs to a degree that would often be scored as ‘no change’ by both CLSI and EUCAST guidelines.^94,95^ Yet in mouse models of gastrointestinal colonization, even therapeutic doses of fluconazole failed to eliminate para-resistant isolates as effectively the non-adapted isolates. This discrepancy suggests that para-resistant cells have increased fitness during drug treatment in a mammalian host as is *in vitro*. Consistent with this model, para-resistant cells evolve resistance to higher drug doses at least an order of magnitude more frequently than the naïve parent. Our findings suggest that the phenotypic plasticity and diversity enabled by this epigenetic state can transmute differences in MIC that would have been systematically ignored by clinical convention into resistance to therapeutic doses of drug in the host.

The strong effects of exposure to subclinical fluconazole doses emphasizes the urgent need to understand the ‘one-health’ consequences of environmental contamination by azole drugs and fungicides as well as other therapeutic agents.^96,97^ These compounds, widely used in medicine and agriculture, are relatively stable; our results identify a mechanism through which their residual levels could broadly impact human, animal, and environmental health.^97^ Furthermore, our work provides compelling evidence that effective treatment of infectious diseases requires not only an understanding of how pathogens respond to drugs, but also how they may adapt over time. Accordingly, it is critical to move beyond simply assessing resistance, which offers a snapshot of the ability to withstand current treatments, to also understand *adaptability*, which predicts potential responses to future treatments. Ultimately, understanding pathogen adaptation to drug exposure can unlock new opportunities to develop therapeutics that either target or avoid the core processes that drive the rapid evolution of drug resistance.

## Acknowledgements

We thank the current and former Jarosz lab members for helpful discussions. We also thank Koon Ho (Chris) Wong and Lakhansing Pardeshi (University of Macau) for assistance with RNA-sequencing analysis. Parts of Fig 1B, Fig 5A, and Fig 5C were created with BioRender.com. J.L.X. is supported by a Stanford University School of Medicine Dean’s Postdoctoral Fellowships, a Mark Foundation Fellow of the Damon Runyon Cancer Research Foundation (DRG 2320–18), and a NIH Pathway to Independence Award (1K99AI169076-01A1). S.Y. holds a Stanford Graduate Fellowship in Science & Engineering and T.S.Y. holds a Stanford Interdisciplinary Graduate Fellowship. L.E. is supported by the Community College Outreach Program at Stanford. K.S.O. is supported by the Crohn’s and Colitis Foundation Career Development Award (DP2AI177827) and the Cifar Azrieli Global Scholars Program. J.B. is supported by a European Research Council Synergy Award (951475 Fungal Tolerance). D.F.J. is supported by the National Institute of Health (DP2-GM119140), the National Science Foundation (NSFMCB116762), and a Vallee Scholar Award. D.F.J. is also a Science and Engineering Fellow of the David and Lucile Packard Foundation.

## Author Contributions

Conceptualization, J.L.X., J.B., D.F.J.; Methodology, J.L.X., S.Y., T.S.Y., K.C., L.H., S.H.K., L.E., C.A.H., N.B., J.R., K.S.O.; Formal Analysis, J.L.X.; Investigation, J.L.X.; Writing – Original Draft, J.L.X., D.F.J.; Writing – Review & Editing, J.L.X., K.S.O., J.B., D.F.J.; Visualization, J.L.X.; Supervision, D.F.J.; Funding Acquisition, J.L.X., D.F.J.

## Methods

### Strains and reagents

All *C. albicans* strains were archived in 25% glycerol and stored at −80 °C. Strains were grown in YPD (1% yeast extract, 2% bactopeptone, and 2% glucose) at 30 °C unless otherwise specified. For solid media, 2% agar was added. Strains were constructed according to standard protocols.^98^ *C. albicans* strains, bacterial plasmids, and oligos used in this study are listed in Supplementary Information (Table S1-3). Fluconazole (TCI Chemicals, F0677-1G) was prepared as a 1 mg/ml stock in water and used at a final concentration of 8 µg/ml for MIC assays, 0.5 µg/ml for the induction of para-resistance, and 0.6 µg/ml in PDR assays. Caspofungin (Sigma-Aldrich, SML0425-5M) was prepared as a 100 µg/ml stock and used at a final concentration of 125 ng/ml in MIC. Phloxine B (Sigma-Aldrich, P2759-25G) was prepared as a 10 mg/ml stock and used at a final concentration of 5 ug/ml in PDR assays.

### Clinical isolates

Respiratory specimens were cultured on blood agar and chocolate agar. Non-respiratory specimens were cultured on BD BBL™ CHROMagar™ Candida (Becton Dickinson). Cultures were incubated at 37 degrees C. Isolates were identified with matrix-assisted laser desorption/ionization-time of flight (MALDI-TOF) mass spectrometry (Bruker).

### Growth curves

Overnight cultures were grown in 3 ml – 5 ml YPD at 30°C. 100 ul of each culture was used to measure OD600 in a 96-well plate (CELLTREAT Scientific, 229596) using a microplate reader (Molecular Devices, SpectraMax ABS Plus). To prepare the inoculum, 1/OD600 x 1.5 µl of the overnight culture was added to 5 ml of YPD. 100 µl of the inoculum (∼10^4^ cells) was added to 100 µl of YPD or 100 µl of YPD with fluconazole at the specified concentration in an edged 96-well plate (Thermo Scientific, 267544). OD600 was measured every 15 min for 48 h in a microplate reader (Biotek, Synergy H1 Hybrid). The generation was calculated using the R package Growthcurver.^99^ The growth curves were plotted in GraphPad Prism (10.2.2).

### Transient drug treatment regime

To assess the effect of transient drug treatment in bulk, inocula were prepared with overnight cultures grown in 3 ml – 5 ml YPD at 30°C. Growth curve experiments were set up in YPD or YPD with fluconazole that inhibits 50% - 80% based on colony forming units (1.63 µM for wild-type SN95) following the steps as described above. After growing for 48 h at 30°C, untreated and fluconazole-treated cultures were used as inocula and the steps for growth curves were repeated.

To assess the effect of drug treatment on individual cells, cultures were grown in YPD or YPD with fluconazole in 4 replicates and diluted 1:50,000 and 1:10,000, respectively. 100 µl of each diluted culture was plated on YPD agar plates (200 – 400 colonies). The plates were incubated at 30°C for 48 hours. 48 single colonies were picked from each condition (12 single colonies from each plate), and each colony was used to inoculate 100 µl of YPD in a 96-well plate. The overnight cultures were mixed and pinned into an edged 96-well plate, each well filled with 100 µl of YPD with 1.63 μM fluconazole, using a 96 long pin repad (Singer Instrument Company, REP-001) with Singer RoToR HDA. Growth was monitored continuously for 48 hours in a microplate reader (Biotek, Synergy H1 Hybrid).

### Selection experiment

For rapid selection of fluconazole and clotrimazole resistance, strains were grown overnight in YPD at 30°C. Cells were plated on YPD or YPD supplemented with 6.03 μM phloxine B and 6.53 μM fluconazole or 742 nM clotrimazole. Plates were left to incubate for 2 days at 30°C in the dark before plates were imaged and quantified visually.

### Phenotypic screen for cross resistance

Each isolate was grown in 200 µl YPD overnight in 96-well plate. The overnight cultures were pinned on solid medium in Singer plates with or without drug in four technical replicates. The colonies were imaged with a scanner after growing for three days at 30°C. For each strain, the average colony size on each drug was normalized for growth on rich medium, and its growth relative to the naïve parent was calculated.

### Permeability-based drug susceptibility assay

For strains with a fluconazole MIC of 1.63 μM, the Permeability-based Drug Susceptibility (PDR) assay plate is made by supplementing YPD agar with 1.96 μM fluconazole and 6.03 μM phloxine B. For streaking: single colonies from YPD were streaked directly onto the PDR plates. For plating: overnight cultures of C. albicans cells were diluted in YPD and 200 – 400 cells were spread onto the PDR plates. The PDR plates were incubated at 30°C for 2 days and imaged face-down on a scanner. Colony number and size were quantified with the Colony Counter plugin in ImageJ (1.54i). Colony numbers were further verified manually. A control was included in every experiment to establish the size and color of colonies that are sensitive to fluconazole. In the presence of fluconazole, sensitive cells accumulate the dye and give rise to small pink colonies, while resistant cells exclude the dye and give rise to large white/lighter pink colonies. Each strain was plated with at least 3 technical replicates and assayed with at least 2 biological replicates. The fluconazole concentration in the PDR plates can be adjusted to correspond to the MIC of the testing strain.

To estimate the induction frequency of para-resistance directly using the PDR plates, cells were either untreated or transiently exposed to fluconazole at MIC and plated on PDR plates. After 2 days of incubation at 30°C, the PDR plates were imaged and the resistant colonies were quantified visually. To confirm the induction frequency, cells were either untreated or transiently exposed to fluconazole at MIC and plated on YPD plates. After 2 days of incubation at 30°C, 96 colonies with no pre-exposure and 288 colonies with pre-exposure to fluconazole were individually added to 100 ul of YPD medium in 96-well plates. After adding 100 ul of 50% glycerol to each well, the cells were mixed, pinned onto YPD Singer plates in 384-well format, and archived in −80°C. The YPD Singer plates were incubated at 30°C for 2 days and cells were subsequently pinned onto YPD and PDR plates. The plates were incubated at 30°C and imaged every 24 hours facing down on a scanner for at least 2 days. The size of the colonies was quantified with SGATool.^100^ The size of the 96 colonies with no prior exposure was used to establish the mean and standard deviation for calculating the z-scores of the 288 colonies with prior exposure to fluconazole. Typically, colony sizes with z-score > 1.5 are considered resistant. To further confirm the colony is para-resistant, the original archive was streaked out on PDR plate. Para-resistant isolates will generally give rise to single colonies that are at least 50% resistant and at least 1% sensitive. However, some para-resistant isolates are unusually stable and only lost after 50+ divisions.

### Whole genome sequencing

Saturated overnight cultures grown in 6 ml of YPD medium were pelleted at 2,190x*g* for 10 min at 4°C. Genomic DNA was extracted using phenol/chloroform as described previously.^41,101^ Sequencing libraries were prepared using the NEBNext® Ultra™ II FS DNA Library Prep with Sample Purification Beads (NEB# E6177). Quality control and quantification were subsequently performed using an Agilent 2100 Bioanalyzer. Samples were submitted for sequencing on the MGI-seq2000 platform at BGI (www.bgi.com/us/home).

Quality control of reads was performed using FastQC (Babraham Institute). Reads were aligned to SC5314 reference genome (Version A21-s02-m09-r10) [van Het Hoog et al., Genome Biology. 2007] using Bowtie2 (Version 2.5.1) [Langmead B and Salzberg S. Nature Methods. 2012]. Mapped reads were sorted and converted using samtools (version 1.17) [Li H et al., Bioinformatics. 2009]. MuTect (Version 1.1.6) [Cibulskis K et al., Nature Biotechnology. 2013] was used to identify mutations in the para-resistant isolates compared to their parent isolates. Sourmash (version 4.8.2) [Brown C and Irber L. JOSS. 2016] was used to create signatures for the a and α alleles and queried against the signatures from raw sequences to identify the presence of the mating alleles.

### RNA sequencing

Overnight cultures were diluted to ∼2×10^5^ cells in 6 ml of YPD with or without 1.63 μM fluconazole. The subcultures were grown until midlog phase at 30°C and cells were harvested by centrifugation at 2,190x*g* for 10 min at 4°C. The pellets were flash-frozen and stored at −80 °C overnight. RNA was isolated using the QIAGEN RNeasy kit. Samples were submitted for sequencing on the BGISEQ-500RS at BGI (www.bgi.com/us/home).

### Mouse gastrointestinal colonization experiments

5-8 week old C57Bl/6 mice (Jackson Laboratory) male and female mice received antibiotic water (0.5 mg/ml ampicillin, neomycin, gentamycin) for 2-7 days prior to oral gavage with 1×10^7 cultured C. albicans (SC5314) and for the duration of the experiment. Antibiotic water was changed weekly. Fluconazole was added to the antibiotic water at 0.5 mg/ml at the indicated experimental time points. All mouse experiments were performed in compliance with federal regulations and guidelines set forth by the University of Utah Institutional Animal Care and Use Committee and the University of Colorado-Anschutz Medical Campus (protocol #01221).

### Minimum inhibitory concentration assay

Antifungal susceptibility was measured in flat-bottom, 96-well plates (CELLTREAT Scientific, 229596) using a broth microdilution protocol described previously.^102^ In brief, minimum inhibitory concentration (MIC) assays were set up in twofold serial dilutions of fluconazole in a final volume of 200 µl YPD per well. Fluconazole gradient was diluted from 26.12 µM to 0.41 µM. Cell densities of overnight cultures were determined in microplate reader (Molecular Devices, SpectraMax ABS Plus) and dilutions were prepared such that ∼10^4^ cells were inoculated into each well. Plates were incubated at 30 °C for 48 hours, at which point the absorbance was determined at 600 nm using a microplate reader and corrected for background from the corresponding medium. Each strain was tested in at least three biological replicates. MIC data was displayed quantitatively with colors using Java TreeView 1.2.0 (http://jtreeview.sourceforge.net).

