## Supplementary Information for "Stress-driven emergence of heritable non-genetic drug resistance"

**Table S1.** ***Candida albicans*** **strains used in this study.**

| Strain Number | Strain Name | Genotype | Source |
|---|---|---|---|
| JBY3532 | SC5314 | Prototrophic | [1] |
| JBY3880 | Chr3 x 3 ABB | SC5314 Chr3 x3 ABB | [2] |
| YDJ4095 | SN95 | *arg4Δ/arg4Δ his1Δ/his1Δ URA3/ura3Δ::imm434 IRO1/iro1::imm434* | [3] |
| YDJ6059 | CaCiS-1 | Prototrophic | This study |
| YDJ6060 | CaCiS-2 | Prototrophic | This study |
| YDJ6065 | CaCiS-7 | Prototrophic | This study |
| YDJ6122 | CaCi-1 | Prototrophic | [4] |
| YDJ6277 | N1 | SN95 naïve lineage 1 | This study |
| YDJ6278 | N2 | SN95 naïve lineage 2 | This study |
| YDJ6279 | N3 | SN95 naïve lineage 3 | This study |
| YDJ6280 | ParaR L1 | SN95 pararesistant lineage 1 | This study |
| YDJ6281 | ParaR L2 | SN95 pararesistant lineage 2 | This study |
| YDJ6282 | ParaR L3 | SN95 pararesistant lineage 3 | This study |
| YDJ6283 | S1 | SN95 sensitive lineage 1 | This study |
| YDJ6284 | S2 | SN95 sensitive lineage 2 | This study |
| YDJ6285 | S3 | SN95 sensitive lineage 3 | This study |
| YDJ6309 | SCUPC2R32A | SC5314 *UPC2A643T-FRT/UPC2* | [5] |
| YDJ9231 | *SNT1* deletion | SN95 *snt1Δ/snt1Δ* | This study |
| YDJ9229 | *SKO1* deletion | SN95 *sko1Δ/sko1Δ* | This study |
| YDJ8230 | *SKO1-GFP* | SN95 *SKO1-GFP-ARG/SKO1* | This study |
| YDJ8522 | *HST1* deletion | SN95 *hst1Δ/hst1Δ* | This study |
| | *MAC1* deletion | SN152 *mac1Δ/mac1Δ* | [6] |
| | *SKO1* deletion | SN152 *sko1Δ/sko1Δ* | [6] |
| | *EFG1* deletion | SN152 *efg1Δ/efg1Δ* | [6] |
| | *ACE2* deletion | SN152 *ace2Δ/ace2Δ* | [6] |
| YDJ9280 | *HOG1* deletion | *SN95 hog1Δ/hog1Δ* | This study |
| YDJ13917 | *CDR1* overexpression | SN95 *TAr-FRT-tetO-CDR1/CDR1* | This study |
| YDJ13959 | *SNT1-GFP* | SN95 *SNT1-GFP-ARG/SNT1* | This study |
| YDJ13981 | *RAP1-GFP* | SN95 *RAP1-GFP-ARG/RAP1* | This study |
| YDJ14006 | ParaR *RAP1-GFP-1* | SN95 *RAP1-GFP-ARG/RAP1* ParaR 4-D1 | This study |

| | | | | |
|---|---|---|---|---|
| YDJ14007 | ParaR *RAP1-GFP-2* | SN95 *RAP1-GFP-ARG/RAP1* ParaR 4-E4 | | This study |

**Strain Constructions**

**YDJ6277/YDJ6278/YDJ6279:** These three isolates were fluconazole-sensitive and had no prior drug exposure. To obtain the single-colony derivatives of the naïve wild-type strain, SN95 was subcultured in YPD at 30°C for 48 hours. Cultures were then plated on YPD agar and incubated at 30°C for an additional 48 hours. Single colonies were subsequently grown in YPD and archived in 25% glycerol. MIC assays confirmed that the archived strains remained fluconazole-sensitive.

**YDJ6280/YDJ6281/YDJ6282:** These three isolates developed resistance following pre-exposure to fluconazole. To obtain the single-colony derivatives, SN95 was subcultured in YPD supplemented with 0.5 µg/ml fluconazole at 30°C for 48 hours. Cultures were then plated on YPD agar and incubated at 30°C for an additional 48 hours. Single colonies were subsequently grown in YPD and archived in 25% glycerol. Growth curves from MIC assays confirmed that the archived strains were adapted to 0.5 µg/ml fluconazole.

**YDJ6283/YDJ6284/YDJ6285:** These three isolates had prior exposure to fluconazole but remained fluconazole-sensitive. To obtain single-colony derivatives following drug pre-exposure, SN95 was subcultured in YPD with 0.5 µg/ml fluconazole at 30°C for 48 hours. Cultures were then plated on YPD agar and incubated at 30°C for an additional 48 hours. Single colonies were subsequently grown in YPD and archived in 25% glycerol. MIC assays confirmed that the archived strains retained fluconazole sensitivity.

**YDJ9231:** PDJ2042 was digested with MssI to release His1-FRT-pENO1-Cas9-pMAL1-FLP-NAT (1 of 2). Fragment A (1066 bp) was amplified from PDJ2045 with oLX191 and oLX192. Fragment B (704 bp) was amplified from YDJ2055 with oLX446 and oLX193. Fragment C (1714 bp) was amplified by stitching Fragment A and Fragment B with oLX199 and oLX200. Donor DNA was generated by annealing oLX447 and oLX448. SNT1 deletion was verified by colony PCR with oLX459 and oLX442 (770 bp for SNT1 deletion and 3774 bp for WT SNT1) and oLX459 and oLX516 (464 bp for WT SNT1, no amplification for deleted allele). The MAL1 promoter was induced to drive expression of FLP recombinase to excise the NAT flipper cassette.

**YDJ9229:** PDJ2042 was digested with MssI to release His1-FRT-pENO1-Cas9-pMAL1-FLP-NAT (1 of 2). Fragment A (1066 bp) was amplified from PDJ2045 with oLX191 and oLX192. Fragment B (704 bp) was amplified from PDJ2055 with oLX510 and oLX193. Fragment C (1714 bp) was amplified by stitching Fragment A and Fragment B with oLX199 and oLX200. Donor DNA was generated by annealing oLX291 and oLX292. SKO1 deletion was verified by colony PCR with oLX300 and oLX277 (430 bp for SKO1 deletion and 2276 bp for WT SKO1) and oLX300 and oLX316 (344 bp for WT SKO1, no amplification for deleted allele). The MAL1 promoter was induced to drive expression of FLP recombinase to excise the NAT flipper cassette.

**YDJ8230:** SKO1-GFP-ARG (3244 bp) was amplified from PDJ1973 with oLX278 and oLX279 and transformed into YDJ4095. Upstream integration (520 bp) was verified with oLX275 and oLX021CaGFP-R. Downstream integration (685 bp) was verified with oLX136 and oLX277.

**YDJ8522:** PDJ2042 was digested with MssI to release His1-FRT-pENO1-Cas9-pMAL1-FLP-NAT (1 of 2). Fragment A (1066 bp) was amplified from YDJ2045 with oLX191 and oLX192. Fragment B (704 bp) was amplified from YDJ2055 with oLX494 and oLX193. Fragment C (1714 bp) was amplified by stitching Fragment A and Fragment B with oLX199 and oLX200. Donor DNA was generated by annealing oLX291 and oLX292. HST1 deletion was verified by colony PCR with oLX497 and oLX498 (390 bp for HST1 deletion and 2358 for WT HST1) and oLX499 and oLX498 (421 bp for WT HST1, no amplification for deleted allele). The MAL1 promoter was induced to drive expression of FLP recombinase to excise the NAT flipper cassette.

**YDJ9280:** PDJ2042 was digested with MssI to release His1-FRT-pENO1-Cas9-pMAL1-FLP-NAT (1 of 2). Fragment A (1066 bp) was amplified from YDJ2045 with oLX191 and oLX192. Fragment B (704 bp) was amplified from YDJ2055 with oLX511 and oLX193. Fragment C (1714 bp) was amplified by stitching Fragment A and Fragment B with oLX199 and oLX200. Donor DNA was generated by annealing oLX298 and oLX299. HOG1 deletion was verified by colony PCR with oLX301 and oLX302 (452 bp for HOG1 deletion and 2373 for WT HOG1) and oLX301 and oLX515 (287 bp for WT HOG1, no amplification for deleted allele). The MAL1 promoter was induced to drive expression of FLP recombinase to excise the NAT flipper cassette.

**YDJ13917:** TAr-NAT-tetO-CDR1 (7175 bp) was amplified from PDJ1763 with oLX312 and oLX313 and transformed into YDJ4095. Upstream integration (433 bp) was verified with oLX317 and oLX314. Downstream integration (397 bp) was verified with oLX315 and oLX318. The NAT cassette and the FLP recombinase were excised by growing the positive colony in YNB-BSA for 2 days at 30C. The SAP2 promoter was induced to drive expression of FLP recombinase to excise the NAT flipper cassette.

**YDJ13959:** SNT1-GFP-ARG (3244 bp) was amplified from PDJ1973 with oLX439 and oLX440 and transformed into YDJ4095. Upstream Integration (486 bp) was verified with oLX441 and oLX128. Downstream Integration (823 bp) was verified with oLX136 and oLX442.

**YDJ13981:** RAP1-GFP-ARG (3244 bp) was amplified from PDJ1973 with oLX433 and oLX434 and transformed into YDJ4095. Upstream integration (490 bp) was verified with oLX435 and oLX128. Downstream integration (390 bp) was verified with oLX136 and oLX436.

**YDJ14006/YDJ14007:** These isolates developed resistance following pre-exposure to fluconazole. To obtain these single-colony derivatives, YDJ13981 was subcultured in 200 µl of YPD supplemented with 0.5 µg/ml fluconazole at 30°C for 48 hours. Cultures were then plated on YPD agar and incubated at 30°C for an additional 48 hours. Single colonies were used to 100 µl of YPD and archived directly in 25% glycerol. A significant increase in colony size on YPD plates supplemented with 0.6 ug/ml fluconazole confirmed that the archived strains were adapted to fluconazole. The strains were subsequently grown in YPD and plated for single colonies on PDR plates. They gave rise to mostly large light colonies and few small dark colonies, confirming that the resistance state is both heritable and reversible.

**Table S2. Bacterial plasmids used in this study.**

| Strain Name | Description | Source |
|---|---|---|
| PDJ2042 | *C. albicans* HIS/FLP CAS9 expression plasmid | [7] |
| PDJ2045 | Universal plasmid template for Fragment A for cloning-free stitching of gRNA expression cassette | [7] |
| PDJ1973 | For tagging genes with GFP at the C-terminus | [8] |

**Table S3 Oligonucleotides used in this study.**

| Strain Name | Description | Source |
|---|---|---|
| oLX128 | GFP+368_R | gtatcaccttcaaacttgac |
| oLX136 | ARG+1161_F | ggagttccatttagagaaac |
| oLX191 | gRNA Universal Fragment A_F | gacggcacggccacgcgtttaaaccgcc |
| oLX192 | gRNA Universal Fragment A_R | caaattaaaaatagtttacgcaag |
| oLX193 | gRNA Fragment B_R | cccgccaggcgctggggtttaaacaccg |
| oLX199 | Fragment C_F | aggtgatgctgaagctattgaag |
| oLX200 | Fragment C for HIS-FLP_R | TAAAGCTGCCACAAGAGGTATTTC |
| oLX277 | SKO1+1994_R | CTTGAAGAAGGTTCAGTAAC |
| oLX291 | CaSKO1 deletion dDNA_F | TCAGATCTTTATTCATTCATTCTTTATTTTTTT TAGTTCCAAAAAGACGGATTGTTCTTGTTCT TGTTATTATTATCATTAAATGTATGTATTTTTG TTT |
| oLX292 | CaSKO1 deletion dDNA_R | AAACAAAAATACATACATTTAATGATAATAAT AACAAGAACAAGAACAATCCGTCTTTTTGGA ACTAAAAAAAATAAAGAATGAATGAATAAAG ATCTGA |
| oLX300 | CaSKO1-172_F | CCACAACATATTCTTCTTGC |
| oLX312 | CaCDR1-M13R_F | TATATCACATATATATTCTATTTATTTTTTGTA CTTAATAATTTCTTTAAAAGGTCAAAAACGAA AAAATTGGAAACAGCTATGACCATG |
| oLX313 | tetO-CaCDR1_R | AAGAAGAGTCTTGACTAATTGCCTTTTCTAAT TTAGATTCATCTTGCGACGACATCTTAGAAT CTGACATcgactatttatatttgtatg |
| oLX314 | CaTAR+72_R | gtacgatggagatagtttac |
| oLX315 | tetO+473_F | atgtttgtcgtttctgatgg |
| oLX316 | CaSKO1+172_R | GCAAATGATCTTTCGAATGG |
| oLX317 | CaCDR1-293_F | TTCTTTTCCACTGGTACTAC |
| oLX318 | CaCDR1+180_R | TGACGAGTCATCTTTGAAAG |
| oLX433 | CaRAP1-GFP_F | TTATAAAGAATATTATGAAAATTGTGAAAAAG TAGGTATGGCTCCTAGATGTTTAACAAGAAT TGCTCCAggtgctggcgcaggtgcttc |
| oLX434 | CaRAP1-ARG_R | TTCATTCTCCCTGATTAACCCTTTAATAATAA AGTTACTTCTTTCTCTGTAAGGTCCTTTCTTT TTTTTTagattaccctgttatcccta |
| oLX435 | CaRAP1+1169_F | CTTGGAGAGATAGATTTAGG |

| | | |
|---|---|---|
| oLX436 | CaRAP1+1610_R | GGGTAAAACTAATGAACGTG |
| oLX439 | CaSNT1-GFP_F | AAAAAATGGTTGGGATGCTATTGTTGCTAATCAGAAAAGTTCGAAAATGTCAATCATGAGTTTGTTAAATggtgctggcgcaggtgcttc |
| oLX440 | CaSNT1-ARG_R | TTGATCTGTTTAACATTATTAACCTATTATTGTTGTTATTTTACTACTGATATATATATCTACATATGTAagattaccctgttatcccta |
| oLX441 | CaSNT1+2886_F | CACAAAATCAACAACCATGG |
| oLX442 | CaSNT1+3410_R | CCCTTGTTTTATTAAGTGGC |
| oLX446 | CaSNT1 gRNA+2310_R | CGTAAACTATTTTTAATTTGGGTTTCACCAGACTCAACAGGTTTTAGAGCTAGAAATAGC |
| oLX447 | CaSNT1 deletion dDNA_F | AACACCATCATTGTCAAATTACTGTCAAAATAATTAATTTTATCATAATGGTACATATGTAGATATATATATCAGTAGTAAAATAACAACAATAATAGGT |
| oLX448 | CaSNT1 deletion dDNA_R | ACCTATTATTGTTGTTATTTTACTACTGATATATATATCTACATATGTACCATTATGATAAAATTAATTATTTTGACAGTAATTTGACAATGATGGTGTT |
| oLX459 | CaSNT1-364_F | TCACTCCTTTGTTGTTGTTG |
| oLX510 | CaSKO1 gRNA+720_R | CGTAAACTATTTTTAATTTGGGGCACTAATAAACTCAGAGGTTTTAGAGCTAGAAATAGC |
| oLX516 | CaSNT1+101_R | GAATAAAAATTTTTCCTTCG |
