## Supplemental Figure for "Stress-driven emergence of heritable non-genetic drug resistance"

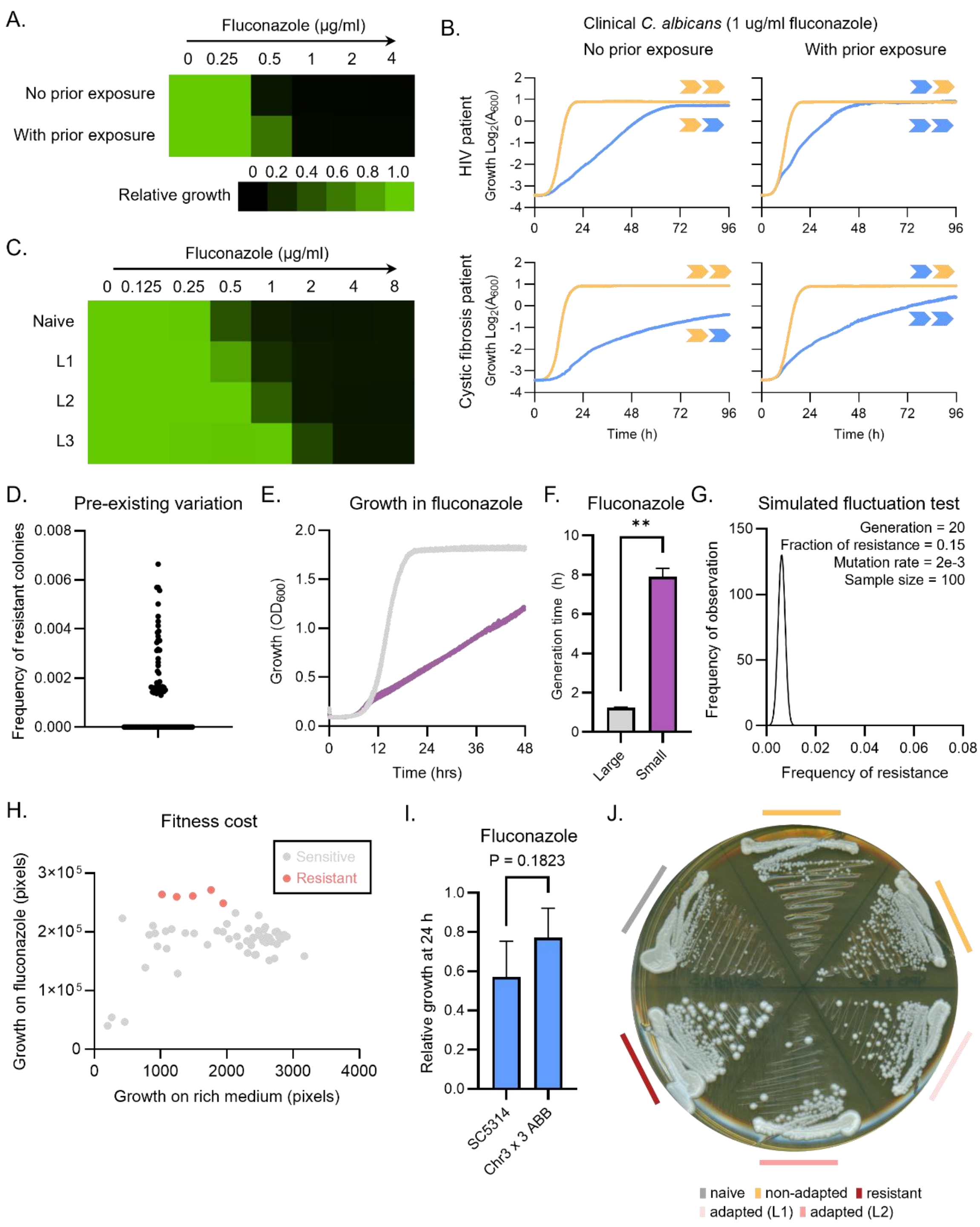


**Figure S1 Characterization of fluconazole adaptation in *C. albicans*.**

(A) Transient exposure to subinhibitory concentrations of fluconazole leads to an increase in MIC. Fluconazole susceptibility assay for SN95 with and without pre-exposure to 1.63 µM fluconazole was conducted in YPD medium. Growth was measured by absorbance at 600 nm after 24 h at 30 °C. Data was quantitatively displayed with color using Treeview (see color bar). (B) Transient exposure to antifungal at near-MIC concentrations primes bulk adaptation to subsequent treatment in two clinical *C. albicans* isolates. Growth of CaCi-1 (top, HIV patient isolate) and CaCiS-2 (bottom, cystic fibrosis patient isolate) in YPD (yellow) and 1.63 µM fluconazole (blue) with no prior exposure to fluconazole (left) and with prior exposure to fluconazole (right). Yellow arrow indicates growth in YPD; blue arrow indicates growth in 1.63 µM fluconazole. (C) The adapted lineages exhibit varying degrees of increase in fluconazole MIC. Growth was measured at 600 nm after 48 h at 30°C. Data was quantitatively displayed with color using Treeview (see color bar in S1A).( D) Estimation of standing variation in fluconazole resistance. Frequency of resistance was measured across 94 replicates. (E) and (F) Large colonies on fluconazole plates are resistant. The growth curves (E) show the growth dynamics and the bar chart (F) shows the generation time of large and small colonies from solid fluconazole medium grown in fluconazole across three biological replicates. Welch's t test, ** $p < 0.01$. (G) Simulation of the fluctuation test assuming 15% of all base substitutions in the *C. albicans* genome could confer dominant resistance to fluconazole. The simulated distribution has a mean of 2.88 and a variance of 3.78 and a variance of 14.29. (H) No anti-correlation between phenotypic resistance to fluconazole and reduced fitness in YPD. Colony size was quantified in ImageJ, comparing between non-adapted (gray) and adapted (salmon) isolates. $r = 0.25$, $p = 0.0544$. (I) An extra copy of Chr3 ABB does not improve growth in fluconazole. Growth at 1.63 µM fluconazole was measured by absorbance at 600 nm after 24 h at 30°C and normalized to the no drug control. Optical densities were averaged across three biological replicates. (J) The adapted lineages produce colonies that vary in size on fluconazole plates. Naïve SN95, non-adapted lineages N1 and N2, adapted lineages L1 and L2, and resistant isolate *UPC2A643T/UPC2* were streaked for single colonies on plates with 1.96 µM fluconazole. Plate was imaged after 48 h at 30°C.

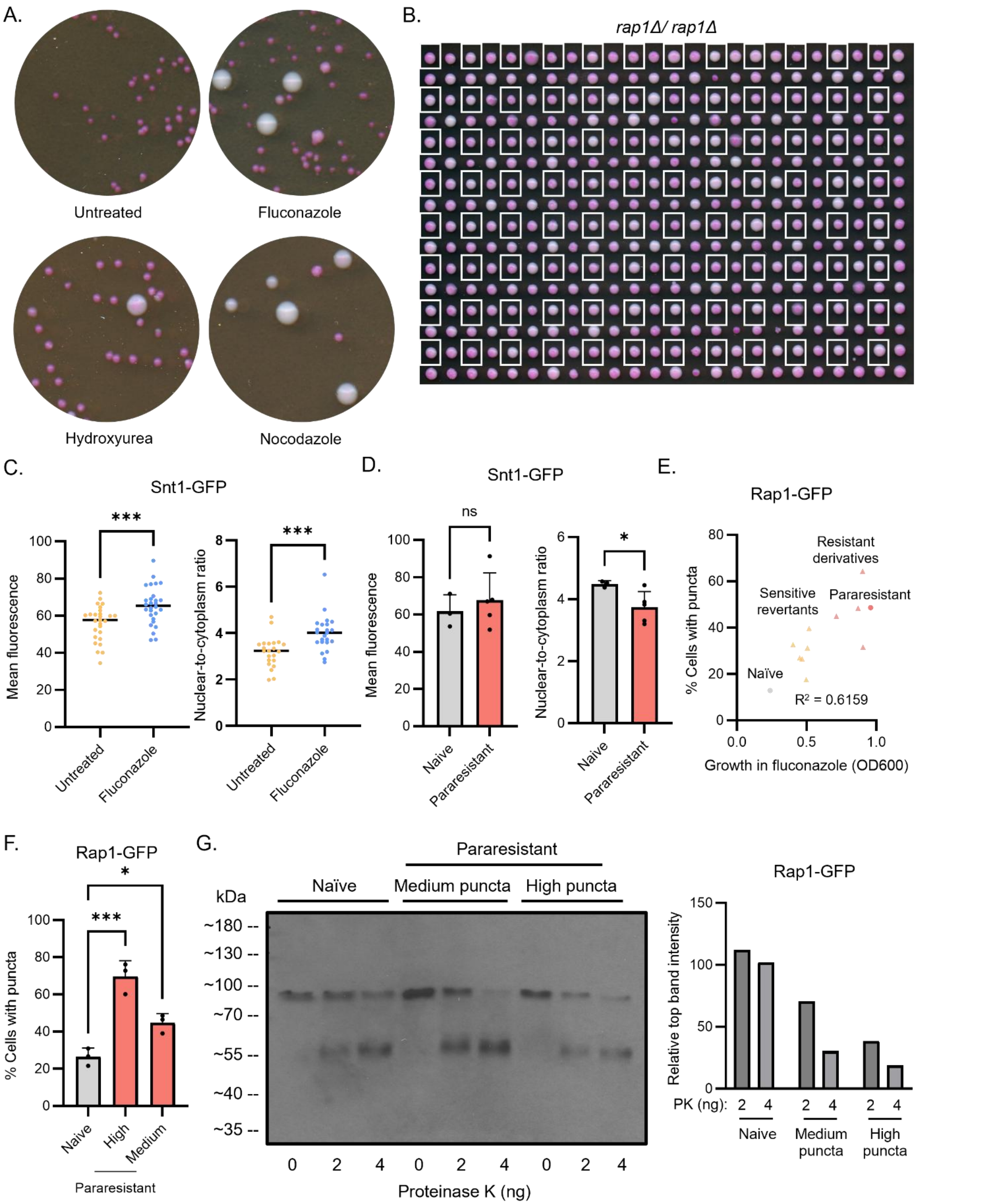


**Figure S2 Para-resistance shares features with [*ESI*⁺].**

(A) Transient exposure to nocodazole but not hydroxyurea induces phenotypic resistance to fluconazole. Wild-type SN95 was transiently exposed to no drug, 1.63 μM fluconazole, 25 μM nocodazole, or 25 mM hydroxyurea, and plated on PDR plates. (B) *RAP1* deletion produces colonies with diverse sizes and hues. Colonies in white boxes were pre-grown in YPD; remaining colonies were pre-exposed to 3.26 μM. Cells were pinned onto YPD before transferred onto PDR plate with 1.96 μM fluconazole. (C) Fluconazole treatment increases Snt1 expression and nuclear accumulation. Naive strain expressing SNT1-GFP was grown in SD+ARG+HIS without (untreated) or with 52.2 μM fluconazole (treated). The mean fluorescence per cell (left) as well as the fluorescence signal in the nucleus and cytoplasm (right) were quantified using ImageJ. Welch's t test, *** $p < 0.001$. (D) Para-resistance affects nuclear distribution but not expression level of Snt1. The mean fluorescence per cell (left) as well as the fluorescence signal in the nucleus and cytoplasm (right) were quantified using ImageJ for naïve (3) and para-resistant (5) isolates. Unpaired one-tailed t test, ** $p < 0.01$. (E) Positive correlation between frequency of Rap1 nuclear puncta and growth in fluconazole. Growth in 3.26 μM fluconazole for naïve control (1), sensitive derivatives (6), resistant derivatives (4), and para-resistant parent (1) were measured by absorbance at 600 nm after 24 h at 30°C. At least 100 cells in 5 fields of view were visually screened and manually counted for nuclear puncta. (F) Para-resistant isolates vary in proportion of cells with nuclear Rap1 puncta. At least 100 cells in 5 fields of view were visually screened and manually counted for nuclear puncta. (G) The pattern of proteinase K degradation for Rap1 is altered in para-resistant isolates. Rap1-GFP was monitored by Western blot and detected with α-GFP antibody (left). Intensity of the top band in samples treated with proteinase K were normalized to the untreated control and quantified using Image J (right).

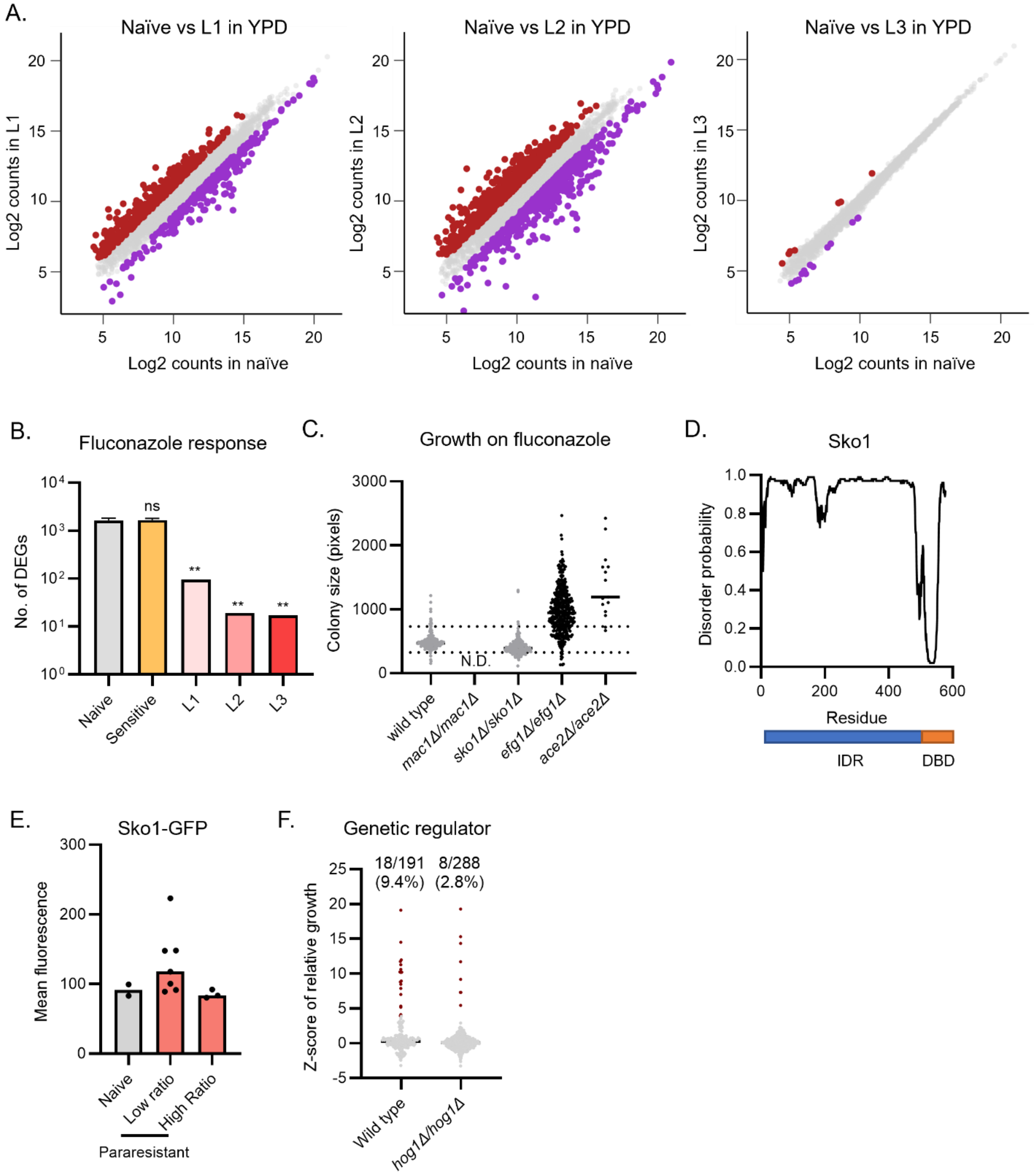


**Figure S3 Chemical, genetic, and transcriptional regulation of para-resistance**

(A) Correlation plots for RNA-seq analysis. Scatterplots highlighting the correlation of normalized RNA-seq read counts for naïve vs L1 (left), naïve vs L2 (middle), and naïve vs L3 (right). Genes upregulated in the para-resistant isolates by 2-fold or more are highlighted in red. Genes downregulated in the para-resistant isolates by 2-fold or more are highlighted in purple. (B) Para-resistant isolates are largely unaffected by fluconazole treatment. The bar plot shows the number of differentially expressed genes between untreated and drug-treated (1.63 μM fluconazole) in naïve isolates (average of N1-N3), sensitive isolates (average of S1-S3) and the three para-resistant lineages (L1-L3). One-way ANOVA, compared to naïve control, ** $p < 0.01$. (C) Deletion of *SKO1* does not affect growth on fluconazole. Wild-type SN152 and transcription factor deletion mutants were plated on PDR plates, and colony sizes were quantified using ImageJ. Dotted line indicates colony sizes that are 1.5-fold smaller or larger than the mean colony size of the wild-type control. (D) Sko1 contains a long stretch of intrinsically disordered region. Disorder probability was predicted by DISOPRED3. (E) The Sko1-GFP expression is similar between naïve and para-resistant isolates. The mean fluorescence per cell was quantified using ImageJ for the same naïve (2) and para-resistant (10) isolates as in Fig 3I. (F) Deletion of *HOG1* blocks the induction of para-resistance. Wild-type SN95 and the *hog1Δ/hog1Δ* mutant were transiently exposed to fluconazole (at 1.63 μM and 0.82 μM, respectively), plated on YPD plates, and single colonies (191 and 288, respectively) were assayed for growth on fluconazole (at 1.96 μM and 0.98 μM, respectively).

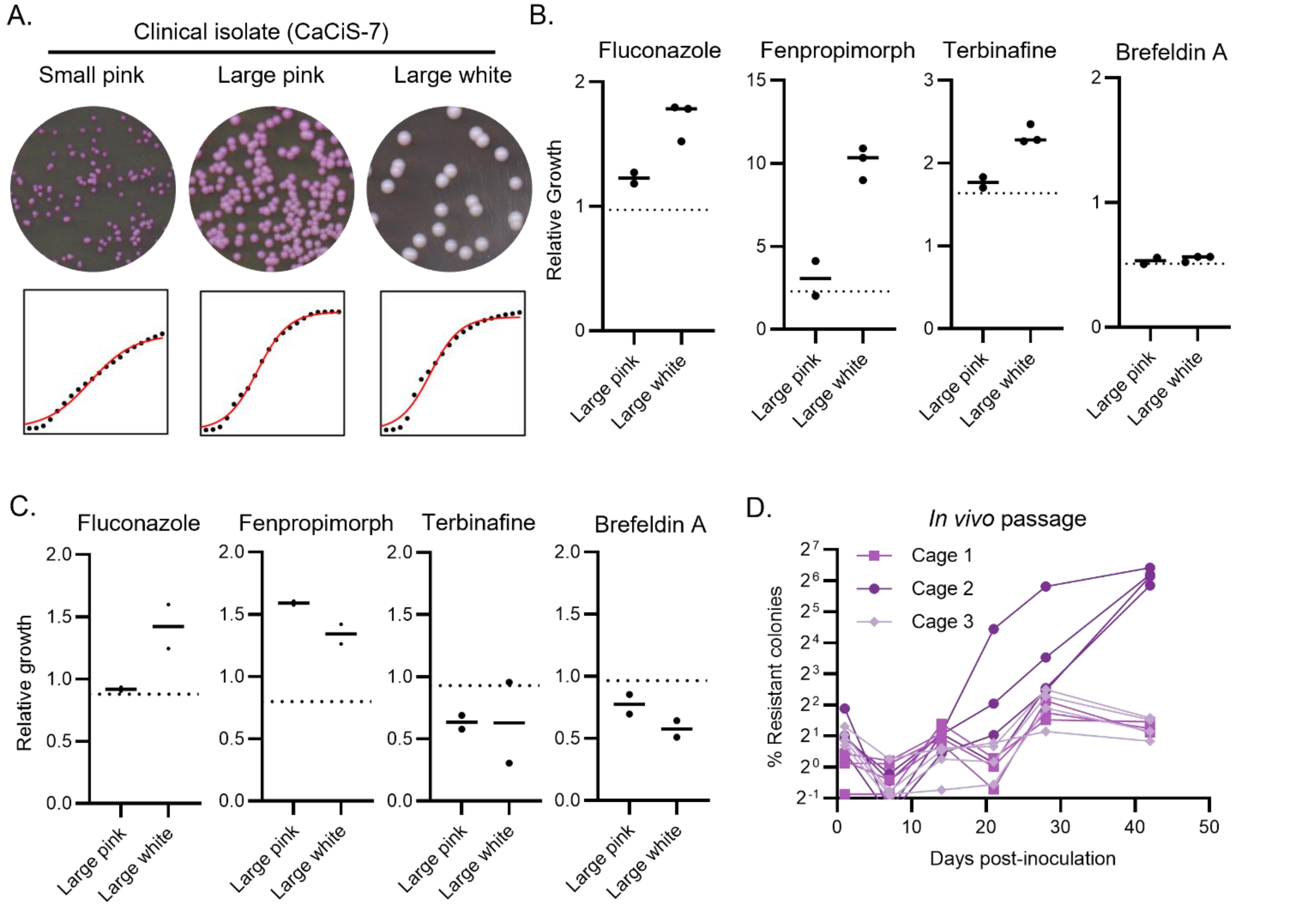


**Figure S4 Phenotypic characterization of *in vivo* isolates of *C. albicans* reveal features of para-resistance.**

(A) Para-resistance in detected in a clinical *C. albicans* isolate. CaCiS-7 (isolated from a cystic fibrosis patient) was plated on YPD plates. 30 single colony derivatives were plated on PDR plates. Growth in 3.62 μM fluconazole was measured over 96 h. (B) CaCiS-1 derivatives exhibit cross-resistance to multiple drugs. The isolates were grown on 1.96 μM fluconazole, 35.7 μM brefeldin A, 34.3 μM terbinafine, and 500 nM fenpropimorph. Colony size was quantified using SGA tools and growth was normalized to the average of small pink derivatives. (C) Derivatives of fecal mouse isolates exhibit cross-resistance to multiple drugs. The isolates were grown on 1.96 μM fluconazole, 35.7 μM brefeldin A, 34.3 μM terbinafine, and 500 nM fenpropimorph. Colony size was quantified using SGA tools and growth was normalized to the average of small pink derivatives. (D) The frequency at which fluconazole resistance emerges *in vivo* varies between cages. Mouse fecal samples were collected every 7 days and plated for single colonies on PDR plates. The plates were imaged after 48 h incubation at 30°C.
